# Four ZIPs contribute to Zn, Fe, Cu and Mn acquisition at the outer root domain

**DOI:** 10.1101/2024.10.31.621270

**Authors:** Kevin Robe, Linnka Lefebvre-Legendre, Fabienne Cleard, Marie Barberon

**Affiliations:** Department of Plant Sciences, University of Geneva, 30 Quai Ernest Ansermet, 1211, Geneva, Switzerland

**Keywords:** ZIP, Zinc, Iron, Copper, Manganese, Transporter, Polarity, Root, Arabidopsis

## Abstract

Zinc (Zn), an essential micronutrient, plays a crucial role in plant development. However, the specific transporters involved in Zn uptake from the soil remain unclear in dicotyledonous. Using promoter-reporter lines in *Arabidopsis thaliana*, we identified several *ZIP* (*Zn-regulated transporter, Iron-regulated transporter (IRT)-like Protein*) family members that are expressed in the epidermis and potentially involved in Zn acquisition from the outer root domain. ZIP2, ZIP3, ZIP5 and ZIP8 predominantly localize to the plasma membrane of epidermal and cortical cells, supporting their potential roles in metal uptake from the soil. Through physiology studies, ionomic profiling and genetic analysis, we determined that ZIP3 and ZIP5 are key contributors to Zn acquisition, while ZIP2 and ZIP8 are primarily involved in copper (Cu) and iron (Fe) acquisition respectively. Notably, ZIP3 and ZIP8 exhibit outer polarity in root epidermal cells, similar to IRT1, underscoring the significance of transporter polarity in mineral acquisition. These findings provide new insights into the mechanisms of metal uptake in plant roots and offer potential strategies for biofortification to enhance metal content in plants.

## Introduction

Plants require at least fourteen essential mineral nutrients for proper growth and development (White et al., 2010). Among these, the transition metal iron (Fe) has been extensively studied, largely due to the pronounced phenotypic consequences of Fe deficiency. In contrast, other transition metals such as manganese (Mn), copper (Cu) and zinc (Zn), while equally critical for various biological processes, have received comparatively less attention. Zinc, in particular, plays a crucial role as a structural and catalytic cofactor in numerous proteins, including RNA polymerase, superoxide dismutase, and alcohol dehydrogenase, and is present in nearly 10% of eukaryotic proteomes (Christine M Palmer & Mary Lou Guerinot, 2009; Sinclair & Krämer, 2012; Stanton et al., 2022). However, in natural soils, Zn is only sparingly soluble, as most of it is bound to minerals such as sphalerite (ZnS), franklinite (ZnFe_2_O_4_), Zn-containing magnetite ([Fe,Zn]Fe_2_O_4_), willemite (Zn_2_SiO_4_), hemimorphite (Zn_4_Si_2_O_7_[OH]_2_ · H_2_O), and zincite (ZnO) (Manceau et al., 2003). Among these, sphalerite is a primary source of soluble Zn^2+^ in the soil solution (Lindsay, 1972). Soil pH is a major factor influencing Zn availability and speciation, with its solubility decreasing as pH increases. Consequently, plants frequently experience Zn deficiency in alkaline soils, which constitute approximately 30% of cultivated land globally (Satoshi Mori, 1999). Both Zn deficiency and excess can result in significant growth defects and yield reduction in the field. To cope with such conditions, plants tightly regulate Zn homeostasis, modulating its uptake, transport, and storage across tissues. Recently, the F group BASIC LEUCINE-ZIPPER transcription factors, AtbZIP19 and AtbZIP23 were identified as intracellular Zn sensors in *Arabidopsis thaliana*. These proteins directly bind Zn^2+^ ions, allowing plants to maintain Zn homeostasis under varying Zn availability (Lilay et al., 2021). This regulatory mechanism for Zn-deficiency response appears to be conserved across species, including rice and in Medicago (Liao et al., 2022; Lilay et al., 2020). Several transporter families play key roles in Zn transport within plants, such as the Zrt-/Irt-related Proteins (ZIP) family, the Yellow Stripe Like (YSL) family, the Heavy Metal ATPase (HMA) family, the Cation Diffusion Facilitator (CDF) family, and the Metal Tolerance Proteins (MTP) family (Christine M Palmer & Mary Lou Guerinot, 2009; Schaaf et al., 2005; Sinclair & Krämer, 2012). In Arabidopsis, for example, Zn efflux into the xylem is primarily mediated by AtHMA2 and AtHMA4, which are predominantly expressed in pericycle cells, while Zn vacuolar storage depends on AtMTP1 (Hanikenne et al., 2008; Kobae et al., 2004; Sinclair et al., 2007). In monocotyledonous plants, Zn can be taken up either as divalent cations (Zn^2+^) or in a chelated form bound to phytosiderophores. However, it has been reported that Zn uptake mediated by ZmYS1 from phytosiderophores in maize is much lower than uptake from ZnSO_4_, suggesting that Zn is primarily acquired as a cation (Von Wirén et al., 1996). Further supporting this, the accumulation of the stable isotope ^67^Zn^2+^ was nearly abolished in the *oszip5oszip9* double mutant of *Oryza Sativa*, indicating that OsZIP5 and OsZIP9 are the primary transporters responsible for Zn acquisition in rice. Both transporters are expressed in the root epidermis and are localized to the plasma membrane (PM) (Huang et al., 2020; Tan et al., 2020; Yang et al., 2020). These findings marked the first demonstration of Zn transporters playing a major role in Zn acquisition in monocotyledonous plants. While the molecular components involved in Zn storage and transport are well characterized, the mechanisms underlying Zn uptake by roots in dicotyledonous plants, including Arabidopsis, remain poorly understood. Several transporters were proposed to facilitate root Zn uptake, including AtIRT1, AtIRT3 and other members of the ZIP family (Lin et al., 2009; Milner et al., 2013; Vert et al., 2002). In Arabidopsis, there are 15 ZIP family members, including 3 IRTs and 12 ZIPs. Yeast strains defective in specific metal transport pathways, such as *zrt1zrt2* for Zn, *fet3fet4* for Fe, *smf1* for Mn or *ctr1* for Cu, were widely used to test ZIP metal transport activity (Grotz et al., 1998; Milner et al., 2013; Rogers et al., 2000; Wintz et al., 2003). ZIP transporters can transport a range of divalent cations, including Zn^2+^, Fe^2+^, Mn^2+^, Cu^2+^, and cadmium (Cd^2+^) (Guerinot, 2000). Although AtIRT3 overexpression results in Zn and Fe accumulation, indicating a possible role in Zn uptake, recent evidence suggest that AtIRT3 primarily functions redundantly in Zn distribution and translocation from root to shoot, without a direct role in Zn uptake from soil (Lee et al., 2021). In contrast, AtIRT1 can transport Zn and other divalent metals in heterologous systems (Connolly et al., 2002; Eide et al., 1996; Rogers et al., 2000; Vert et al., 2002), but its role *in planta* for Zn uptake appears to be restricted to specific conditions, such as Fe deficiency when its expression is induced. Moreover, recent findings show that Fe deficiency-induced Zn accumulation is independent of AtIRT1 (Quintana et al., 2022). Several other ZIPs, including AtZIP1, AtZIP2, AtZIP3, AtZIP4, AtZIP5, AtZIP6, AtZIP7, AtZIP9, AtZIP11, and AtZIP12 were shown to complement yeast mutants defective in Zn uptake, suggesting that they may contribute to Zn transport in plants (Grotz et al., 1998; Milner et al., 2013; Nguyen et al., 2022). While many studies have reported Zn transport in heterologous systems, *in planta* characterization of ZIP transporters remains limited. For instance, Wu et al., 2009 observed a slight increase in shoot Zn content in the *zip5* mutant compared to the wild type (WT), but no significant difference was detected in the *zip6* mutant. The *zip2* mutant exhibited Zn overaccumulation in roots, while *zip1* showed no difference in Zn content relative to WT (Milner et al., 2013). In contrast, recent findings reported reduced Zn translocation from root to shoot in the quadruple *irt3zip4zip6zip9* mutant (Lee et al., 2021). Promoter activity analysis suggests that these four ZIP transporters are involved in Zn transport to the stele, but not directly in Zn uptake from the soil.

In our search for transporters involved in Zn uptake in the dicotyledonous Arabidopsis, we first assessed the expression of the entire *ZIP* family in roots under both Zn-sufficient and Zn-deficient conditions and investigated their promoter activity in roots. Based on these data, we selected ZIP transporters expressed at the root periphery as putative candidates for Zn acquisition and characterized their functional roles. Our *in planta* evidence, derived from mutants and overexpressing lines, indicate that ZIP2 and ZIP8 are primarily involved in Cu and Fe acquisition, respectively, while ZIP3 and ZIP5 are involved in Zn acquisition. Consistent with their functions, ZIP2, ZIP3, ZIP5 and ZIP8 localize at the PM and endomembrane compartments of epidermal and cortical cells. The polarity of epidermal ZIP3 and ZIP8 towards the outer root domain (facing the soil) emphasizes their significance in Zn and Fe acquisition from the soil and underscores the importance of transporter polarity in their functional roles. Overall, this study highlights the value of re-evaluating transporter localization to uncover new functions. The identification of ZIP3 and ZIP5 as key transporters involved in Zn acquisition from the soil marks a significant advancement in our understanding of mineral nutrition, as Zn was one of the last essential elements for which transporters involved in acquisition had not been unequivocally identified in dicotyledonous plants.

## Results

### Physiological effect of zinc deficiency in *Arabidopsis thaliana*

To investigate the effect of Zn deficiency on plant physiology, we cultivated WT plants in zinc-sufficient (15 µM Zn; +Zn) and deficient (0 µM Zn; –Zn) media. After three weeks of growth on horizontal plates under –Zn conditions, the plants stunted growth and chlorosis, resulting in a significant reduction in both fresh weight (FW) and chlorophyll content compared to those grown in +Zn conditions (Figure 1A-C). As anticipated, the diminished chlorophyll content correlated with a decrease in the maximum quantum yield of photosystem II (PSII, F_v_/F_m_), which reflects the potential efficiency of PSII, further confirming the impact of Zn-deficiency on photosynthetic performance (Figure 1D). While mineral deficiencies often induce significant compensatory changes at the whole plant level, the specific effect of Zn deficiency on root and shoot ionome profiles have not been extensively documented. To address this gap, we conducted an ICP-OES (Inductively Coupled Plasma – Optical Emission Spectrometry) analysis of mineral content in the roots and shoots of 3-week-old plants grown in +Zn or –Zn conditions. As expected, Zn was nearly undetectable in both roots and shoots of WT plants under –Zn conditions (Figure 1E, 1F). Moreover, we observed significant reductions in the content of potassium (K), magnesium (Mg), Mn, sodium (Na), strontium (Sr), calcium (Ca), and Cu in the shoots of plants grown in –Zn compared to those grown in +Zn. Conversely, the levels of Fe, molybdenum (Mo) and aluminum (Al) remained largely unaffected by Zn deficiency in shoots. Notably, Zn deficient roots accumulated more than three time more Mn than control roots, suggesting that the transport systems involved in Zn uptake may also facilitate Mn uptake under Zn-deficient conditions. Additionally, we found that Mg, Sr, Al, Fe and K levels were significantly reduced in the roots under –Zn conditions while the contents of Ca, Cu, Mo and Na remained unchanged. We also assessed the translocation of minerals from roots to shoots under both +Zn and –Zn conditions. The root-to-shoot translocation of Al, Ca, Cu, K, Mn, Mo, Na and Sr decreased under –Zn condition. In contrast, the translocation of Fe and Zn increased under –Zn conditions. The translocation of Mg from roots to shoots remain consistent between +Zn and –Zn conditions (Figure S1A). Overall, these findings establish conditions under which Zn deficiency significantly affects plant development, physiology and overall mineral accumulation. Furthermore, they suggest that compensatory mechanisms may be at play for other minerals in –Zn conditions.

**Figure 1.**
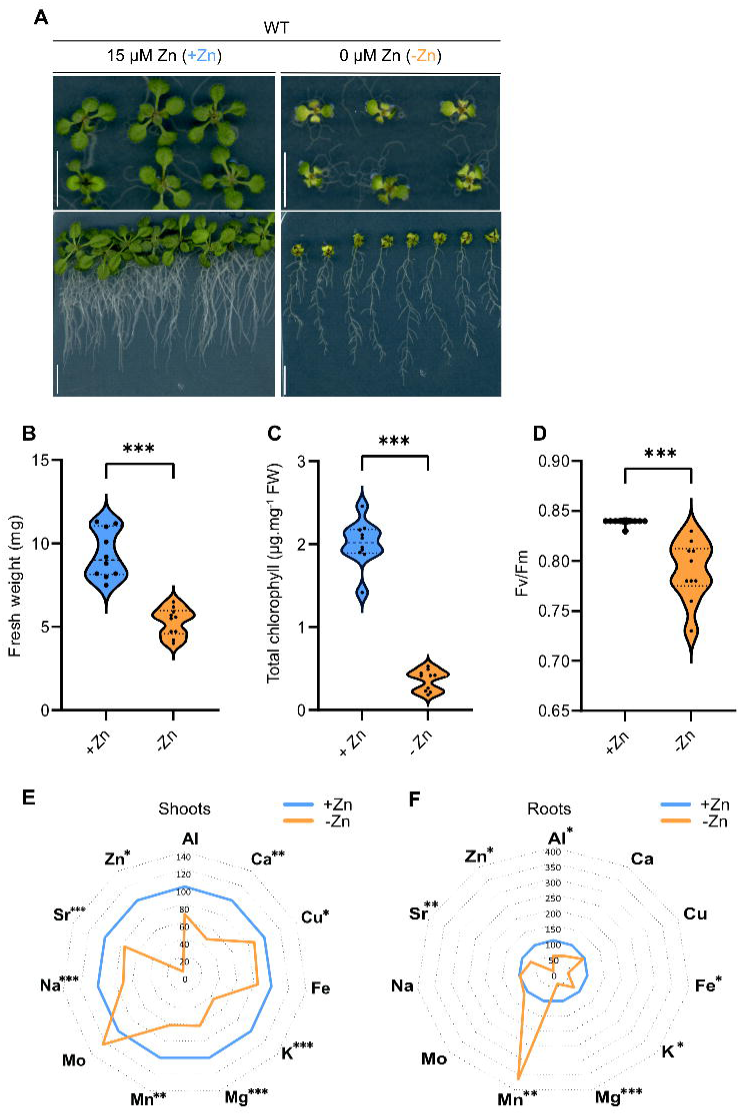
Impact of Zinc deficiency on plants development and ionome. (**A**) Representative images of WT plants grown for 3 weeks in Zn-sufficient (+Zn) and Zn-deficient conditions (-Zn), displayed on horizontal (top) and vertical plates (bottom). Scale bars: 1 cm. Violin plots showing **(B)** fresh weight (FW), **(C)** chlorophyll content, and **(D)** Fv/Fm of dark-adapted WT plants grown 3 weeks in Zn-sufficient (+Zn, blue) or Zn-deficient conditions (-Zn, orange) conditions. In the violin plots dashed lines represent the median and dotted lines represent the first and third quartiles (n≥10) **(E)** Shoots and **(F)** roots ionome profiles of WT plants grown for 3 weeks under Zn-sufficient (+Zn) or Zn-deficient (-Zn) conditions. Data are represented relative to +Zn conditions, set at 100, and correspond to the average values from 3 independent pools, each consisting of at least 10 individual shoots and roots. Numerical values are presented in Table S1. **B-F** Statistical differences were determined using Student’s *t*-test or Mann Whitney test, (\**P* < 0.05; \*\**P* < 0.01; \*\*\**P* < 0.0001).

### Spatio-temporal expression analysis of the *AtZIP* family reveals potential candidates for Zn acquisition

To gain insight into the transcriptional response of plants under Zn-deficient conditions and identify candidates for Zn uptake from the soil, we first assessed the expression of the entire *ZIP* family in WT plants grown for 1 week in either +Zn or –Zn conditions (Figure 2A). The expressions of *IRT1*, *ZIP4*, *ZIP7*, *ZIP10* and *ZIP11* were not significantly altered by Zn deficiency (Figure 2B, Figure S1B). In contrast, we observed increased mRNA accumulation of *IRT3*, *ZIP1*, *ZIP3*, *ZIP5*, *ZIP6, ZIP9* and *ZIP12* in WT roots grown under Zn deficiency compared to control conditions, while *IRT2, ZIP2* and *ZIP8* showed decreased expression. These results suggest that the upregulated transporters may serve as potential candidates for Zn uptake in conditions of limited Zn availability. Notably, *ZIP2* was the only member of the *ZIP* family that exhibited high expression under control conditions, indicating its potential role in basal metal uptake (Figure S1B). This observation aligns with previous reports of high *ZIP2* expression in control conditions (Milner et al., 2013). Anticipating that transporters involved in Zn uptake from the soil would be expressed at the root periphery, we next examined the spatial expression pattern of the 15 ZIP transporters using promoter-reporter lines. We investigated promoter activity in the root tip (zone I), the fully elongated zone (zone II) and the differentiated root (zone III) in control conditions. The promoters were fused to express a fluorescent reporter, NLS-3xmVenus in 5-day-old seedlings (Figure 2C). Promoters of *IRT1*, *IRT2*, *ZIP2*, *ZIP3*, *ZIP5* and *ZIP8* displayed activity primarily at the root periphery in zone II and III (*i.e.* mainly in epidermal and cortical tissues), while the promoter of *ZIP7* was predominantly active in central root tissues such as the endodermis and the stele. Z*IP11* also showed activity in central root tissues but was mostly active in cortical cells. *IRT3*, *ZIP1* and *ZIP10* exhibited almost exclusively endodermal promoter activity (with *ZIP10* being active only in zone III), while *ZIP6* promoter activity was primarily detected in the pericycle in both zones II and III. Promoter activity for *ZIP4*, *ZIP9* and *ZIP12* was minimal under our experimental conditions, with *ZIP12* displaying extremely low signals at the root periphery and in the inner tissue of the zone I. These observations suggest that ZIP2, ZIP3, ZIP5 and ZIP8 may be involved in metal uptake from the soil at the root periphery, similarly to the well-characterized IRT1. Meanwhile, IRT3, ZIP1 and ZIP10 could play a role in metal uptake and transport within the endodermis. Interestingly, six *ZIP* promoters (*IRT2*, *IRT3*, *ZIP1*, *ZIP2*, *ZIP3* and *ZIP7*) showed activity in cells at the root periphery in zone I, indicating their potential role in mediating the transport of divalent transition metal ions in the root tip. Given that the root tip continuously forages the soil for nutrients and water, it is plausible that these ZIP transporters may also serve a sensing function, akin to IRT1(Robe & Barberon, 2023). Based on our expression analysis and promoter reporter lines, we selected ZIP2, ZIP3, ZIP5 and ZIP8 for further characterization as potential candidates for metal uptake from the soil. ZIP2 was of particular interest due to its outer promoter activity, its strong expression in control conditions, and its ability to complement *ctr1* and *smf1* yeast strains. Given these features, we hypothesize that ZIP2 may play a role not in Zn uptake, but in the acquisition of Cu and Mn from the soil (Figure 2B, 2C and Figure S1B, S1C, Milner et al., 2013; Wintz et al., 2003). ZIP8, on the other hand, was investigated as a candidate for Fe acquisition due to its specific epidermal promoter activity, increased expression under Fe deficiency, and its close homology to IRT1 (Figure 2A, 2C, Figure S1D). ZIP3 and ZIP5 were selected as prime candidates for Zn uptake from the soil based on multiple lines of evidence. Both genes were strongly upregulated under Zn deficiency (Figure 2B and Figure S1B) and are known to be regulated by the transcription factors *bZIP19* and *bZIP23* (Assunção et al., 2010). Additionally, their promoter activity was primarily detected at the root periphery (Figure 2C), and both have been shown to complement the yeast *zrt1zrt2* strain, further supporting their role in Zn acquisition (Milner et al., 2013; Nguyen et al., 2022).

**Figure 2.**
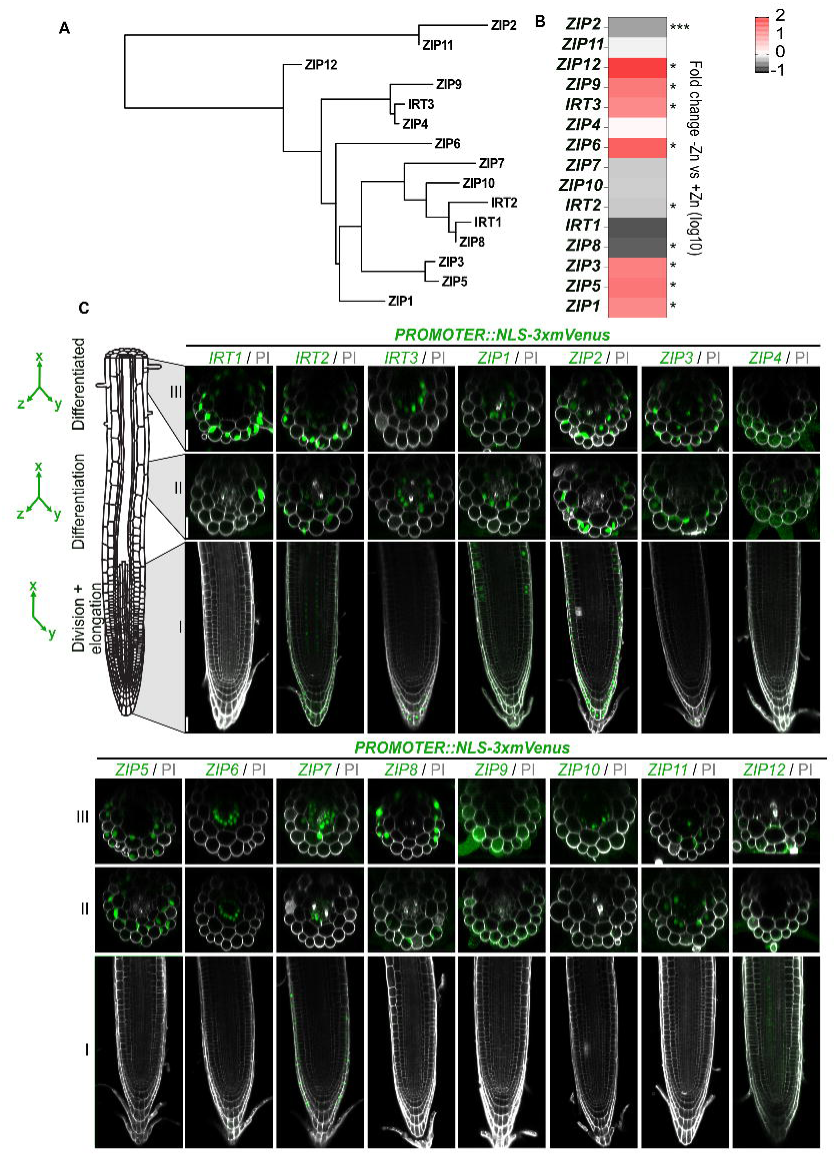
Spatio-temporal expression of the *AtZIP* family in roots. **(A**) Phylogenetic tree of the 15 members of AtZIP family in *Arabidopsis thaliana*. The tree was generated using protein sequences with Phylogeny.fr. **(B)** Relative transcript levels of the fifteen *ZIP* members in WT plant roots grown 1 week under control conditions (+Zn) or Zn-deficient conditions (-Zn). Results are presented as a heatmap, highlighting fold changes compared to control conditions (+Zn). Numeric values are provided in Figure S1B. Data represent 4 biological replicates, with each replicate being a pool of at least 40 roots. Statistical differences were determined using Student’s *t*-test or Mann Whitney test with significance levels indicated (\**P* < 0.05; \*\**P* < 0.01; \*\*\**P* < 0.0001). **(C)** Representative images of *ZIP::NLS-3xmVenus* expression pattern in different root zones: zone I (root tip, lower pictures), zone II (early differentiated zone, middle pictures), and zone III (mature zone, upper pictures). Nuclear-localized mVenus signal is shown in green while PI used to stain the cell wall is shown gray. Images were captured with identical settings for each line along the root, but settings varied between lines due to differences in promoter activity among *ZIPs*. For zone I, a singe focal plant for mVenus and PI is displayed (XY), while for zone II and III, mVenus signal is presented as a maximum projection of Z-stacks (XYZ), overlaid with a single orthogonal view of PI extracted from the Z-stacks. Images correspond to T1 plants; at least 5 independent T1 plants were imaged. The signal was confirmed in T2 plants. Scale bars: 25 µm for all images.

### ZIP2, ZIP3, ZIP5 and ZIP8 localize to the epidermis-soil interface

To confirm the potential role of ZIP2, ZIP3, ZIP5 and ZIP8 in metal acquisition from the soil, we first examined their localization in roots. Transgenic lines were generated expressing a fusion of the *ZIP* genomic DNA with the coding sequence for mCitrine (mCit), under the control of their endogenous promoters, within their respective loss-of-function mutant backgrounds to ensure functional complementation (see below). Based on previous studies that demonstrated the successful integration of mCitrine in the second extracellular loop of IRT1 while maintaining its functionality (Dubeaux et al., 2018), we applied the same strategy for ZIP2/3/5/8-mCitrine constructs. Structural modeling of the fusion proteins predicted that inserting mCitrine into these ZIPs transporters did not alter their 3D conformation, similar to IRT1-mCitrine (Figure S2). Since we expected these transporters to localize at the PM under certain conditions, we co-localized the ZIP transporters with FM4-64, a marker for the PM and endocytic system.

In control, Cu-or Mn-deficient conditions, ZIP2-mCitrine predominantly accumulated in the epidermis of root zone II and in both epidermal and cortical cells of the differentiated root zone III (Figure 3A). Some signal was also detected in the vasculature (white arrows). ZIP2-mCitrine showed partial PM localization under control conditions but was also present in intracellular structures, which co-localized with FM4-64 after 10 minutes, suggesting its localization in endodermal compartments (Figure S3A, S3B). This dual PM-endosomal localization is common for PM proteins fused to highly sensitive fluorophores, possibly reflecting intermediate trafficking stages from the endoplasmic reticulum (ER) to the PM, as observed for transporters like IRT1, NRAMP1 (NATURAL RESISTANCE-ASSOCIATED MACROPHAGE PROTEIN 1) and PHO1 (PHOSPHATE 1) (Barberon et al., 2011; Castaings et al., 2021; Vetal & Poirier, 2023). For IRT1, such localization also reflects substrate-controlled cycling between the PM and early endosomes/Tran-Golgi Network (EE/TGN) (Barberon et al., 2011; Dubeaux et al., 2018; Spielmann et al., 2022). In ZIP2, Cu and Mn deficiency did not notably alter cell-specific localization, except for the near absence of ZIP2-mCitrine in the stele under Cu-deficient conditions (Figure 3A). Considering that transporter polarity can affect their function, we evaluated the potential polarity of ZIP2 in the elongation zone using *UBQ10::ZIP2-mCitrine* overexpression line (Figure 3B). This line was used because ZIP2 expression under its endogenous promoter was undetectable in the elongation zone, where polarity can be more accurately assessed. By calculating the ratio of outer to inner PM fluorescence, we found ZIP2 to be apolar in the epidermis, with a fluorescence ratio close to 1 (Figure 3C).

**Figure 3.**
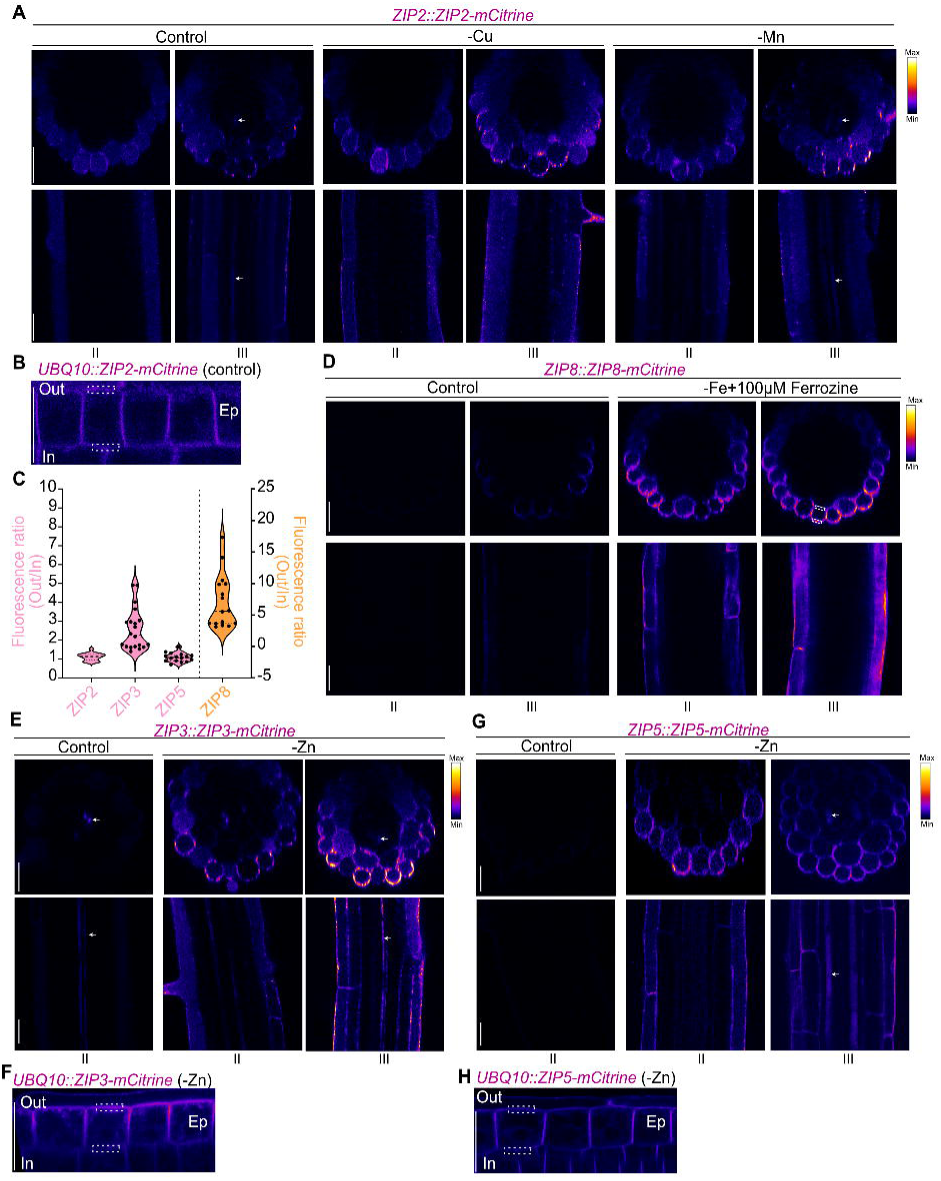
Localization of ZIP2, ZIP8, ZIP3 and ZIP5 in the outer root domain. (**A**, **D**, **E** and **G**) Representative confocal images showing mCitrine fluorescence in 5-day-old seedlings expressing ZIP2, ZIP8, ZIP3 or ZIP5 fused to mCitrine under the control of their endogenous promoters. Seedlings were grown in either control or metal-deficient conditions (-Cu, –Mn, –Fe + 100 µM Ferrozine, –Zn). The upper panels display root transversal views (extracted from Z-stacks), while the lower panels show longitudinal views taken at the median focal plane of the root. The corresponding root zone (II or III, as defined in Figure 2C) is indicated below the images. All images were acquired from T2 plants. (**A**, **B**, **D-H**) mCitrine fluorescence is displayed using a Fire look-up-table (LUT). **(B, F, H)** Confocal images of mCitrine fluorescent signal in 5-day-old seedlings expressing *UBQ10::ZIP2/3/5-mCitrine* grown under control (ZIP2) or –Zn conditions (ZIP3 and ZIP5). Roots were imaged in the median focal plane of epidermal (Ep) cells within the elongation zone to assess polar localization. **(C)** Fluorescence ratio analysis for ZIP2, ZIP3, and ZIP5-mCitrine shown in panels **B, F** and **H** (pink). The ZIP8-mCitrine ratio (orange) was calculated in the epidermis of root zone III. The fluorescence ratio was determined by dividing the average fluorescence intensity at the outer (Out) PM by the intensity at the inner (In) PM for each epidermal cell. The same region of interest (ROI) was used for both outer and inner fluorescence quantification. An example of the ROI used to quantify Out/In ratio is shown with the dashed square for each genotype (**Figure B, D, F, H**). Ratios were calculated from at least 5 cells per root and at least 3 independent roots. The dashed lines in the violin plots represent the median, while the dotted lines represent the first and third quartiles (n≥15). Scale bars: 25 µm for all images.

Next, we investigated ZIP8 localization, given its potential role in Fe acquisition. Under, control conditions, ZIP8-mCitrine was barely detectable in root zone II but was observed at low levels in the epidermis of root zone III (Figure 3D). Under Fe deficiency (-Fe + 100 µM Ferrozine), ZIP8-mCitrine strongly accumulated at the PM of the epidermal cells in both zone II and III (Figure 3D, S3C). Interestingly, ZIP8-mcitrine displayed pronounced outer polar localization in the epidermis under both control and Fe-deficient conditions, reinforcing its possible involvement in metal acquisition from the soil (Figure 3C, S3C).

We also examined the localization of ZIP3 and ZIP5. In control conditions, ZIP3-mCitrine was detected in a few cells in the vasculature (white arrows, Figure 3E). Under Zn-deficient conditions, ZIP3-mCitrine accumulation significantly increased, with prominent expression in the epidermis and cortex, in addition to the vasculature (white arrows). ZIP3-mCitrine predominantly localized to the PM in the elongation zone, though subcellular localization was also observed, co-localizing with FM4-64 indicating these structures were part of the endocytic system (Figure S3D, S3E). Due to the low accumulation of ZIP3-mCitrine in the root tip, we assessed ZIP polarity in the elongation zone using the overexpression line, revealing outer polar localization (facing the soil) in the epidermis under Zn deficiency (Figure 3C, 3F). Lastly, ZIP5-mCitrine localization was explored. In control conditions, ZIP5-mCitrine was undetectable in roots, but it accumulated in the epidermis of root zone II under Zn deficiency (Figure 3G). In zone III, ZIP5-mCitrine accumulated in the epidermis, cortex and some cells in the vasculature under Zn-deficient conditions (white arrows, Figure 3G). ZIP5-mCitrine predominantly localized to the PM but unlike ZIP3, did not show polar localization in the epidermis (Figure S3F, S3G, 3C, 3H).

### ZIP2 mediates Cu acquisition

Given ZIP2’s expression at the root periphery and its PM localization in epidermal cells under Cu and Mn deficiency, as well as its ability to complement *smf1* and *ctr1* yeast strains, we hypothesized that ZIP2 could facilitate Cu and Mn acquisition from the soil. To investigate this, we first isolated the previously described T-DNA insertion line *zip2-1*, which exhibits no detectable *ZIP2* expression (Figure S4A, S4B, Milner et al., 2013), and generated a new CRISPR/Cas9-induced allele, *zip2-2_cr_* (Figure S4A). The *zip2-2_cr_* mutant harbors a 68 bp deletion resulting in a premature stop codon (Figure S4C). When grown in soil, *zip2-2_cr_* mutants exhibited a slight increase in shoot weight compared to both WT and *zip2-1* (Figure S4D, S4E). However, when plants were grown on agar plates containing 0.05 µM Cu (+Cu), shoot FW, root length and chlorophyll content were similar across all genotypes (WT, *zip2-1,* and *zip2-2_cr_*) (Figure 4A-D). In contrast, when Cu was omitted from the growth medium (-Cu), both *zip2-1* and *zip2-2_cr_* exhibited significantly reduced shoot FW and root length compared to WT plants (Figure 4A-D). This increased sensitivity to Cu deficiency in the *zip2* mutants suggests a critical role for ZIP2 in Cu acquisition. To explore whether ZIP2 might also mediate Mn uptake, we grew *zip2* mutants and WT plants on Mn-deficient plates. However, unlike under Cu deficiency, neither *zip2-1* nor *zip2-2_cr_* displayed notable differences compared to WT under Mn-deficient condition (-Mn) (Figure S4F-H). The lack of a pronounced phenotype suggests that ZIP2 plays no or a limited role in Mn uptake. To further confirm ZIP2’s role in Cu and Mn acquisition, we measured Fe, Zn, Cu and Mn content in both shoots and roots of *zip2* mutants grown for 3 weeks on control agar plates. These metals were selected as they represent the primary substrates for ZIP transporters. Notably, both *zip2-1* and *zip2-2_cr_* mutants accumulated significantly less Cu than WT in both roots and shoots, reinforcing the idea that ZIP2 plays a major role in Cu acquisition (Figure 4E, S4I). While shoot concentrations of other metals remained unchanged in the *zip2* mutants, a slight reduction in root Mn content was observed in both mutants (Figure 4E and S4I), though this reduction was less pronounced compared to Cu. Given the strong reduction in Cu content in *zip2-1*, we used this phenotype to assess the functionality of the ZIP2-mCitrine fusion protein used in localization studies (Figure 3A). Complementation of Cu and Mn content in roots of two independent *zip2-1/ZIP2::ZIP2-mCitrine* lines confirmed the functionality of the fusion protein and the causal role of ZIP2 in these phenotypes (Figure 3A, S4J). Overall, our data strongly suggest that ZIP2 primarily mediates Cu acquisition from the soil, with a potential, though more limited, role in Mn acquisition.

**Figure 4.**
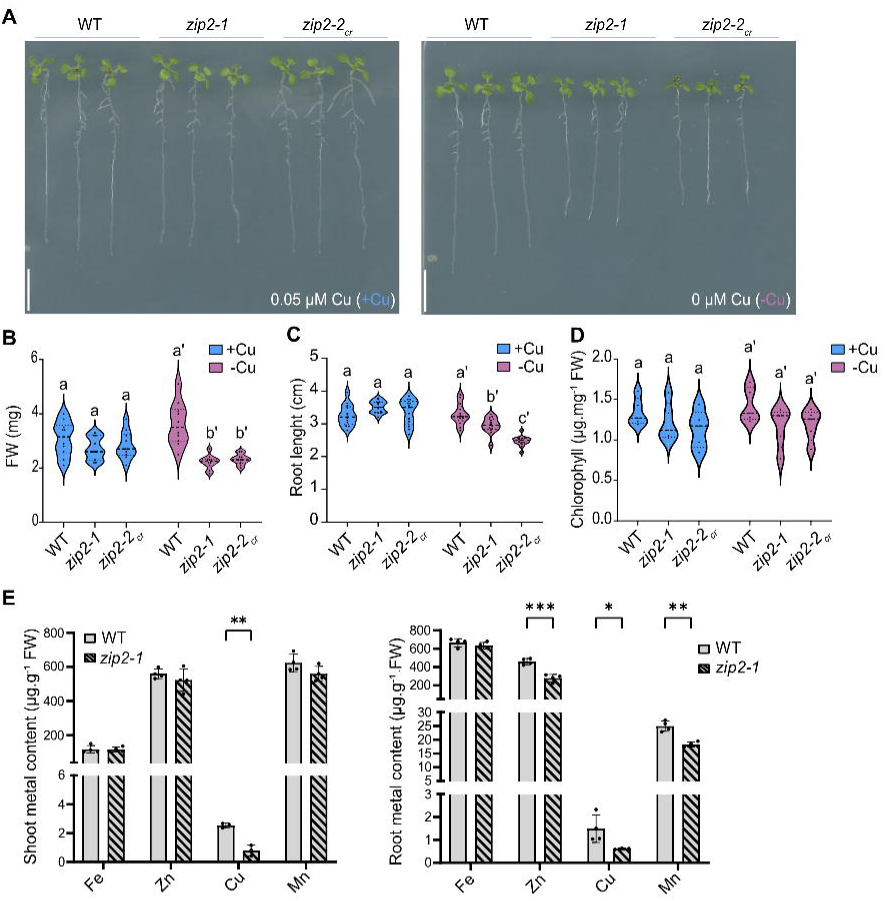
Impaired Cu accumulation in the *zip2* mutant. **(A)** Representative images of WT and two *zip2* mutant alleles grown vertically for 10 days under control conditions (+Cu) and copper deficient conditions (-Cu). Three representative plants for each genotype were transferred to agar plates for imaging. Scale bars: 1 cm. (**B**-**D)** Violin plots showing the distribution of **(B)** fresh weight (FW), **(C)** root length, and **(D)** chlorophyll content of plants grown in +Cu or –Cu for 10 days (n≥15 for B and C, n≥5 for D). Dashed lines of the violin plots represent the median, while dotted lines mark the first and third quartiles. Different letters indicate statistically significant differences between genotypes, as determined by one-way ANOVA followed by Tukey’s post hoc test or Kruskal–Wallis test followed by Dunn’s test (*P* < 0.05). **(E)** Metal content (Fe, Cu, Mn, Zn) in shoots (left) and roots (right) of WT and *zip2-1* mutant grown vertically for 3 weeks on control agar plates. For each biological replicate, 3-4 plants were pooled (n=4). Data are represented as mean ± SE, with significant differences between genotypes determined by Student’s *t*-test or Mann-Whitney test (\**P* < 0.05; \*\**P* < 0.01; \*\*\**P* < 0.0001).

### ZIP8 mediates Fe acquisition

Given ZIP8’s protein localization and expression patterns (Figure 3D, S1D), we hypothesized that ZIP8 may participate in Fe uptake from the soil, similarly to its closest homologue, IRT1 (Figure 2A). As no T-DNA insertion line were available for the 5’ coding region of *ZIP8*, we generated two CRISPR/Cas9-induced alleles. The *zip8-1_cr_* allele carries a deletion in the promoter and the 5’ coding sequence, predicted to produce a truncated 2 aa peptide (Figure S5A, S5B), while *zip8-2_cr_* harbors a single A insertion in the 5’ coding sequence, leading to a premature stop codon (Figure S5A, S5C). We began by investigating the phenotype of WT *zip8-1_cr_*, *zip8-2_cr_*, and *irt1-2* plants grown on soil for 3 weeks. Since IRT1 is the only known ZIP transporter involved in Fe uptake, we used *irt1-2* as a reference mutant. Unlike *irt1-2*, which exhibited strong chlorosis, neither of the *zip8* mutants showed obvious chlorotic symptoms when grown on soil (Figure S5D). However, both *zip8* alleles displayed a slight reduction in shoot FW compared to WT (Figure S5E). Next, we grew WT and the two *zip8* mutants for 10 days on control (+Fe, 50 µM Fe) and Fe-deficient (-Fe, 0 µM Fe) agar media to assess their sensitivity to Fe deficiency. We analyzed shoot FW, root length and chlorophyll content, as indicators of Fe deficiency response. However, no significant difference in these traits were observed between WT and *zip8* mutants under either Fe condition (Figure 5A-D), suggesting that ZIP8 does not play a major role in Fe deficiency. To further explore ZIP8’s role in metal acquisition, we measured metal content (Fe, Cu, Mn, and Zn) in shoots and roots of WT and *zip8* mutants grown on control agar plates for 3 weeks. While Mn and Zn levels were comparable between WT and *zip8* mutant in roots, Fe and Cu content in the roots of *zip8* alleles were significantly reduced compared to WT (Figure 5E). Interestingly, no significant changes in metal content were observed in the shoots of *zip8* mutants, suggesting that ZIP8 primarily mediates metal acquisition in the outer root tissues without substantially affecting root-to-shoot translocation. To confirm the functionality of the ZIP8-mCitrine fusion protein, we used the *UBQ10::ZIP8-mCitrine* reporter line. This line showed increased Zn and Mn content, indicating that the fusion protein used for protein localization was functional (Figure 3D, Figure S3C, S5F). Similarly, overexpression of *IRT1* increased Zn and Mn levels in roots, without significantly altering Fe content (Figure S5G). In conclusion, our results demonstrate that ZIP8 is involved in Fe acquisition in the outer root tissues and, similar to IRT1, is involved in the transport of other divalent ions such as Cu^2+^, Mn^2+^, and Zn^2+^. However, ZIP8’s role in Fe transport appears to be more specialized and does not fully overlap with IRT1’s function.

**Figure 5.**
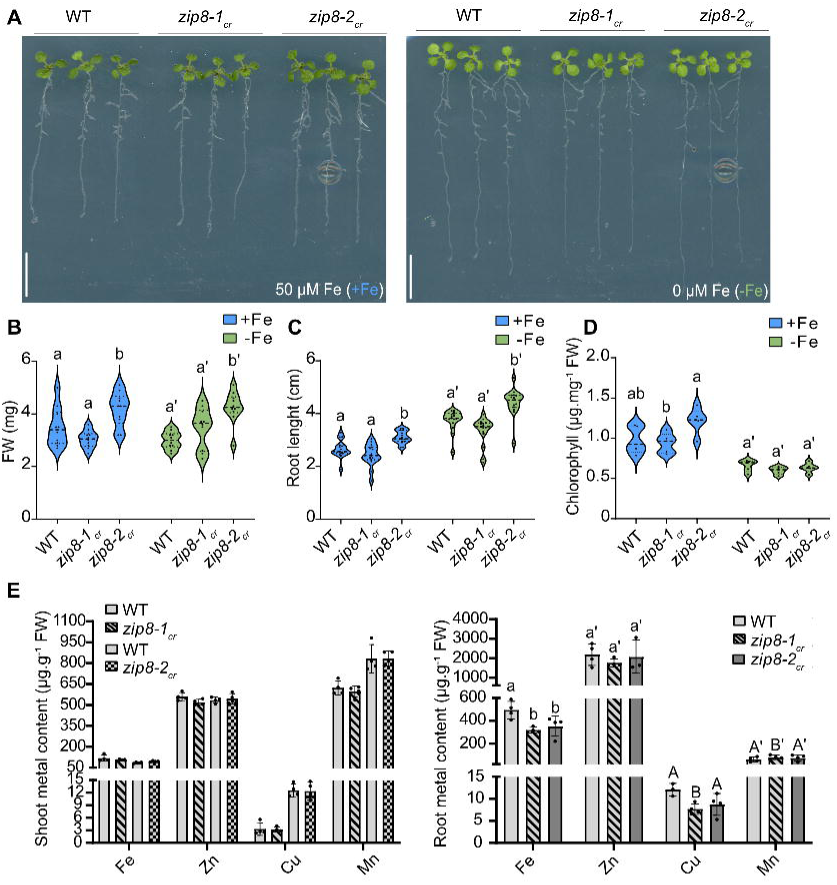
Impaired Fe accumulation in *zip8* mutant. **(A)** Representative images of WT and two *zip8* mutant alleles grown vertically for 10 days under control conditions (+Fe) and Fe-deficient conditions (-Fe). Three plants per genotype were transferred onto an agar plate for imaging. Scale bars: 1cm. (**B**-**D)** Violin plots showing the distribution of **(B)** FW, **(C)** root length, and **(D)** chlorophyll content of plants grown as described in panel A (n≥15 for B and C, n≥5 for D). Dashed lines of the violin plots represent the median, while dotted lines mark the first and third quartiles. Different letters indicate statistically significant differences between genotypes, as determined by one-way ANOVA followed by Tukey’s post hoc test or Kruskal–Wallis test followed by Dunn’s test (*P* < 0.05). **(E)** Metal content (Fe, Cu, Mn, Zn) in shoots (left) and roots (right) of WT and *zip8* mutants grown vertically for 3 weeks on control conditions. For each biological replicate, 3-4 plants were pooled (n=4). Data are represented as mean ± SE, with significant differences between genotypes determined by Student’s *t*-test or Mann-Whitney test for shoot data where no significant differences were observed, while root data were analyzed using one-way ANOVA followed by Tukey post hoc test (*P* < 0.05).

### ZIP3 and ZIP5 mediate Zn acquisition

Given the localization of ZIP3 and ZIP5 at the root periphery under Zn-deficient conditions (Figure 3E, 3G), along with prior evidence from yeast *zrt1zrt2* complementation assays, we hypothesized that these transporters could be involved in Zn uptake from the soil. Due to the close phylogenic relationship between ZIP3 and ZIP5 (Figure 2A), we generated a double mutant (*zip3-1zip5-1*) by crossing the single KO T-DNA insertion mutants *zip3-1* and *zip5-1* to account for potential functional redundancy (Figure S6A). Additionally, we created CRISPR/Cas9 mutants for *ZIP3* and *ZIP5* (*zip3-2_cr_* and *zip5-2_cr_*, Figure S6B, S6C). The *zip3-2_cr_* mutant carries a substitution of two CC nucleotides with a single A, resulting in a premature stop codon, while the *zip5-2_cr_* mutant has a single T insertion that also introduces a premature stop codon (Figure S6B, S6C). We first assessed the growth of WT, *zip3-1, zip5-1, zip3-1zip5-1* and the CRISPR-induced mutants *zip3-2_cr_* and *zip5-2_cr_* on soil. No significant differences in FW were observed between WT and the mutants (Figure S6D, S6E). When grown on agar plates with sufficient Zn (15 µM Zn, Control), the *zip3-1zip5-1* double mutant exhibited larger overall growth compared to WT and the single mutants, but no significant differences in root length or chlorophyll content were noted (Figure 6A-D). Under Zn-deficient conditions (-Zn), WT plants and the single mutants displayed increased root length, consistent with previous findings (Niehs et al., 2024). Strikingly, the double mutant had significantly smaller shoots and reduced chlorophyll content compared to WT under Zn deficiency, indicating heightened sensitivity to low Zn (Figure 6B-D). The single mutants showed little to no phenotype, suggesting that ZIP3 and ZIP5 function redundantly in Zn acquisition. To further support the role of ZIP3 and ZIP5 in Zn acquisition, we measured metals content in roots and shoots of 3-week-old plants grown on agar plates. While Fe, Zn, Cu and Mn levels in shoots were similar between WT and *zip3-1zip5-1*, Zn levels in roots were significantly reduced in the double mutant, confirming that ZIP3 and ZIP5 are involved in Zn acquisition (Figure 6E). Metal analysis of soil-grown plants also revealed lower Zn (and, to a lesser extent, Cu) levels in *zip3-1zip5-1* compared to WT, whereas Fe and Mn levels remained unaffected, indicating that ZIP3 and ZIP5 may also influence Zn accumulation under conditions involving evapotranspiration (Figure S6F). Given the known role of transpiration in mineral translocation (Muro et al., 2023), and the accumulation of ZIP3 and ZIP5 in the vasculature, they may contribute to Zn distribution to sink tissues. However, the similar root-to-shoot Zn translocation ratios between WT and *zip3-1zip5-1* suggest that their contribution to Zn partitioning is limited and that other transporters also are likely involved (Figure S6G, Lee et al., 2021). To dissect the individual contributions of ZIP3 and ZIP5, we examined metal content in the single mutants. Soil-grown shoot analysis revealed that ZIP3 is primary responsible for the Zn deficiency observed in *zip3-1zip5-1,* as *zip5-1* and *zip5-2_cr_* mutants accumulated Zn at levels comparable to WT (Figure S7A-D). In contrast to *zip3* mutant, no significant differences in root Zn content were detected between WT and the *zip5* mutant, suggesting that ZIP5 plays a more limited role in Zn (Figure S7E, S7F). However, elevated Zn levels in the *zip3-1/ZIP3::ZIP3-mCitrine* and in *zip5-1/ZIP5::ZIP5-mCitrine* complementation lines confirmed the functionality of the fusions and the causality of the *zip3* and *zip5* phenotypes (Figure 3E, 3H, S7B, S7F). To gain further insight into the distribution of Zn in the *zip3-1zip5-1* mutant, we used the Zinpyr-1 fluorescent probe (Sinclair et al., 2007). In WT differentiated roots, Zinpyr-1fluorescence was primarily localized in the stele and endodermis, indicating Zn accumulation in these tissues. In contrast, fluorescence intensity was significantly reduced in the *zip3-1zip5-1* double mutant, though the overall pattern of distribution remained unchanged (Figure 6F, 6G). Metal measurements and Zinpyr-1 staining in *ZIP3* and *ZIP5* overexpression lines confirmed that the overexpression increased Zn accumulation across root cell layers (Figure 6H, 6I), with no overaccumulation of other metals typically transported by ZIP family members (e.g. Cu, Mn, Fe). Overall, these data indicate that ZIP3 and ZIP5 are redundantly involved in Zn acquisition from the soil in Arabidopsis, with ZIP3 playing a more dominant role. The fact that *irt1* mutants accumulate WT-like Zn levels in several studies, raises questions about the extent of IRT1’s contribution to Zn uptake in soil-grown plants (Barberon et al., 2011; Quintana et al., 2022). To address this, we employed a promoter-swap approach, expressing *IRT1* under *ZIP2*, *ZIP3* and *ZIP5* promoters (Figure S7G). Plants expressing *ZIP2::IRT1* and *ZIP3::IRT1* in the *zip3-1zip5-1* background restored Zn accumulation to WT levels in shoots, confirming IRT1’s capacity for Zn transport *in planta* (Figure S7H). Overexpression of IRT1 (*35S::IRT1*) further corroborated this finding (Figure S5F). However, *ZIP5::IRT1* failed to rescue Zn levels in *zip3-1zip5-1* shoots, likely due to the lack of *ZIP5* promoter activity in the vasculature under our conditions (Figure 2C). Overall, these results suggest that Zn acquisition predominantly depends on ZIP3 activity at the root periphery.

**Figure 6.**
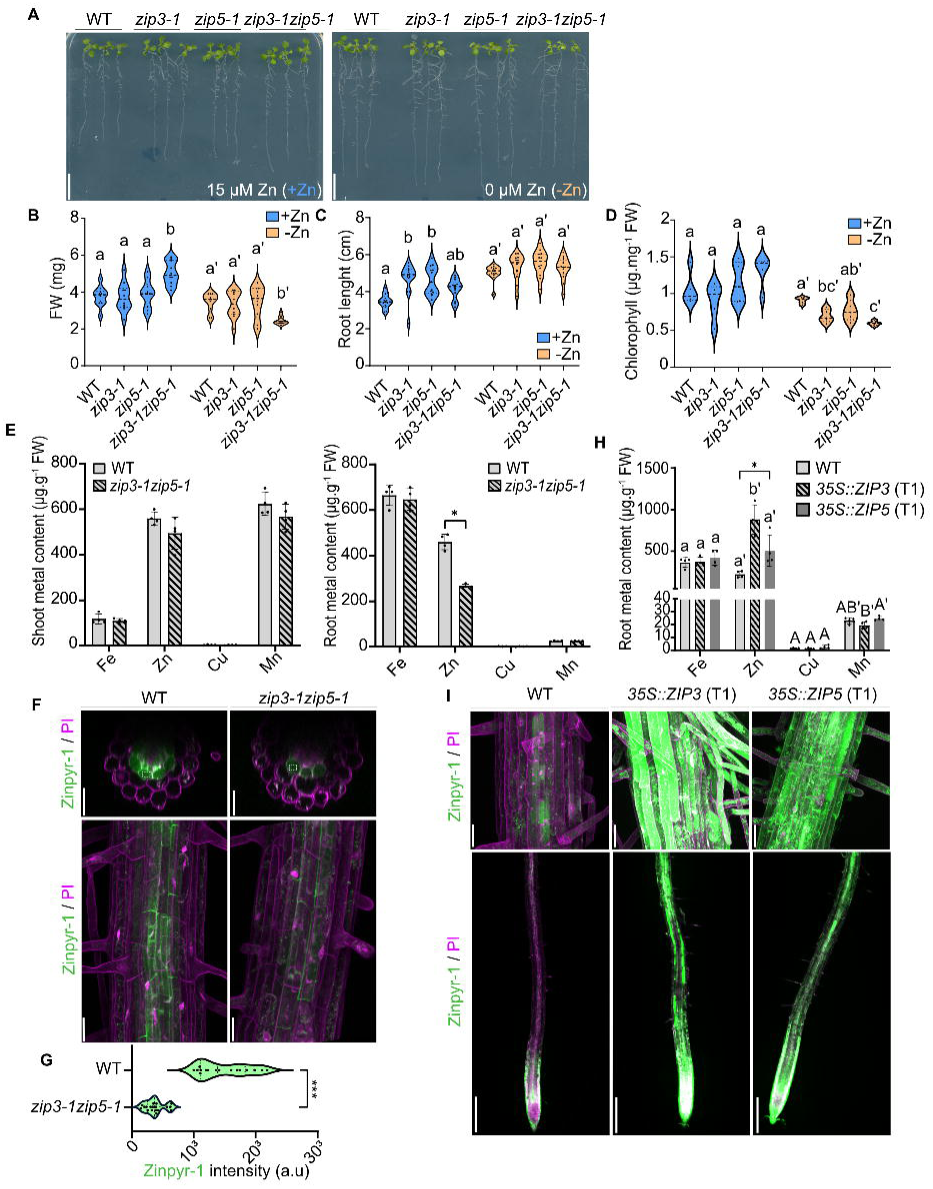
Zn uptake by ZIP3 and ZIP5. **(A)** Representative images of WT plants, *zip3-1*, *zip5-1* and the double mutant *zip3-1zip5-1* grown vertically for 10 days under control conditions (+Zn) and Zn-deficient conditions (-Zn). Three representative plants from each genotype were transferred to an agar plate for imaging. Scale bars: 1cm. (**B**-**D)** Violin plots showing the distribution of **(B)** FW, **(C)** root length, and **(D)** chlorophyll content of plants grown as described in panel A (n≥15 for B and C, n≥5 for D). Dashed lines of the violin plots represent the median, while dotted lines mark the first and third quartiles. Different letters indicate statistically significant differences between genotypes, as determined by one-way ANOVA followed by Tukey’s post hoc test or Kruskal–Wallis test followed by Dunn’s test (*P* < 0.05). **(E)** Metal content (Fe, Cu, Mn, Zn) in shoots (left) and roots (right) of WT and *zip3-1zip5-1* mutants grown for 3 weeks on control agar plates. For each biological replicate, 3-4 plants were pooled (n=4). Data are represented as mean ± SE, with significant differences between genotypes determined by Student’s *t*-test or Mann-Whitney test (\**P* < 0.05; \*\**P* < 0.01; \*\*\**P* < 0.0001). **(F)**. Representative confocal images of 5-day-old WT and *zip3-1zip5-1* roots labeled with 20 µM Zinpyr-1 for 3 h. Plants were grown under Zn-deficient conditions for 5 days prior to staining. Upper panels show cross-sectional views and the lower panels show maximum projections of the differentiated root. Scale bars: 25µm. **(G)** Violin plots showing the distribution of endodermal Zinpyr-1 signal quantified in region of interest (ROI) in WT and *zip3-1zip5-1* roots. At least four independent roots were analyzed (n≥15). In the violin plots dashed lines represent the median and dotted lines represent the first and third quartiles. Statistical difference between genotypes was determined using Mann-Whitney test (\**P* < 0.05; \*\**P* < 0.01; \*\*\**P* < 0.0001). **(H)** Metal content (Fe, Cu, Mn, Zn) in roots of WT, ZIP3 and ZIP5 overexpression lines grown for 3 weeks on control agar plates. For each biological replicate, 3-4 T1 plants were pooled (n=4). Data are presented as mean ± SE, with different letters indicating statistically significant differences between genotypes, determined by one-way ANOVA followed by Tukey’s post hoc test (*P* < 0.05). **(I)** Representative confocal images of 5-day-old WT, *35S::ZIP3* (T1) and *35S::ZIP5* (T1) plants stained with 20 µM Zinpyr-1 for 3 h. Plants were grown for 5 days on 1 µM Zn prior to staining. Upper panels show a cross section of the differentiated root (scale bars: 25 µm), and lower panels show maximum projections from the root tip (scale bars: 200 µm).

### Zn excess triggers ZIP3 endocytosis

Nutrient transporters are frequently regulated at multiple levels, including transcriptional, post transcriptional and post translational mechanisms, in response to nutrient availability (Barberon et al., 2011; Dubeaux et al., 2018; Takano et al., 2005, 2006; Tanaka et al., 2011; Vert et al., 2002). To investigate if Zn availability influences ZIP3 stability in the root, we exposed *UBQ10::ZIP3-mCitrine* plants to 150 µM Zn, a 10-fold excess relative to standard conditions (Lešková et al., 2017). By using the *UBQ10* promoter, we bypassed potential transcriptional regulation of *ZIP3* to focus on post-transcriptional effects. ZIP3 was chosen because of its predominant role in Zn acquisition from the soil. Confocal microscopy revealed that, despite the Zn excess, the overall ZIP3-mCitrine fluorescence did not significantly decrease compared to Zn-deficient condition (Figure 7A, 7B). This suggests that ZIP3 protein abundance is not markedly affected by Zn levels, or that higher Zn concentrations might be required to influence its stability. Although ZIP3-mCitrine accumulation was unchanged, we hypothesized that Zn availability might impact its subcellular localization, as it is well-established for other transporters that undergo endocytosis in response to excess substrates (Castaings et al., 2021; Dubeaux et al., 2018; Ivanov & Vert, 2021; Takano et al., 2005). Previous colocalization with FM4-64 indicated that ZIP3 is present both at the PM and within the endocytic system ( Figure S3E). We examined ZIP3 localization in the root tip, an optimal region for subcellular compartment imaging due to its small cell size. For endosomes detection, we imaged the surface focal plane and, for assessing ZIP3 polarity, we focused on the median plane of epidermal cells. Plants were grown in the presence of 15µM Zn for 5 days, then exposed to either Zn deficiency (0 µM Zn, –Zn) or a 10-fold Zn excess (150 µM Zn, 10xZn) for 16 hours. Under –Zn conditions, ZIP3-mCitrine predominantly localized to the PM, displaying outer polarization, as described earlier (Figure 3C, 3F, 7C). However, upon exposure to excess Zn, the intracellular ZIP3 signal increased while PM localization diminished (Figure 7C, 7D). Quantification of the PM-to-intracellular (PM/Intra) ratio of ZIP3-mCitrine revealed a significant reduction in this ratio, indicating delocalization of ZIP3 in response to Zn excess (Figure 7E). To determine if this intracellular signal was due to ZIP3 endocytosis upon Zn excess, we treated the plants with Brefeldin A (BFA), a fungal toxin that inhibits vesicle recycling between endosomes and PM. Since BFA also blocks the secretion of newly synthesized PM proteins, we pre-treated plants with cycloheximide (CHX) to inhibit protein synthesis, allowing us to track only the pool of already translated ZIP3-mCitrine. Under –Zn conditions, ZIP3-mCitrine was found at the PM as well as in BFA-induced intracellular bodies, suggesting that ZIP3 cycles between the PM and endocytic compartments under Zn deficiency (Figure 7F). Following Zn excess, the majority of the ZIP3-mCitrine signal accumulates in BFA bodies, as indicated by the significantly altered PM/Intra ratio (Figure 7F, 7G, S8A). This confirms that ZIP3 undergoes rapid endocytosis in response to Zn excess, likely as a protective mechanism to prevent excessive Zn acquisition as previously shown for IRT1 (Barberon et al., 2011; Dubeaux et al., 2018; Spielmann et al., 2022). Whether ZIP3 is subsequently targeted for vacuolar degradation remains an open question. Interestingly, unlike IRT1 whose localization is regulated by its secondary substrates, ZIP3 localization appears to be modulated by the availability of its primary substrate, Zn.

**Figure 7.**
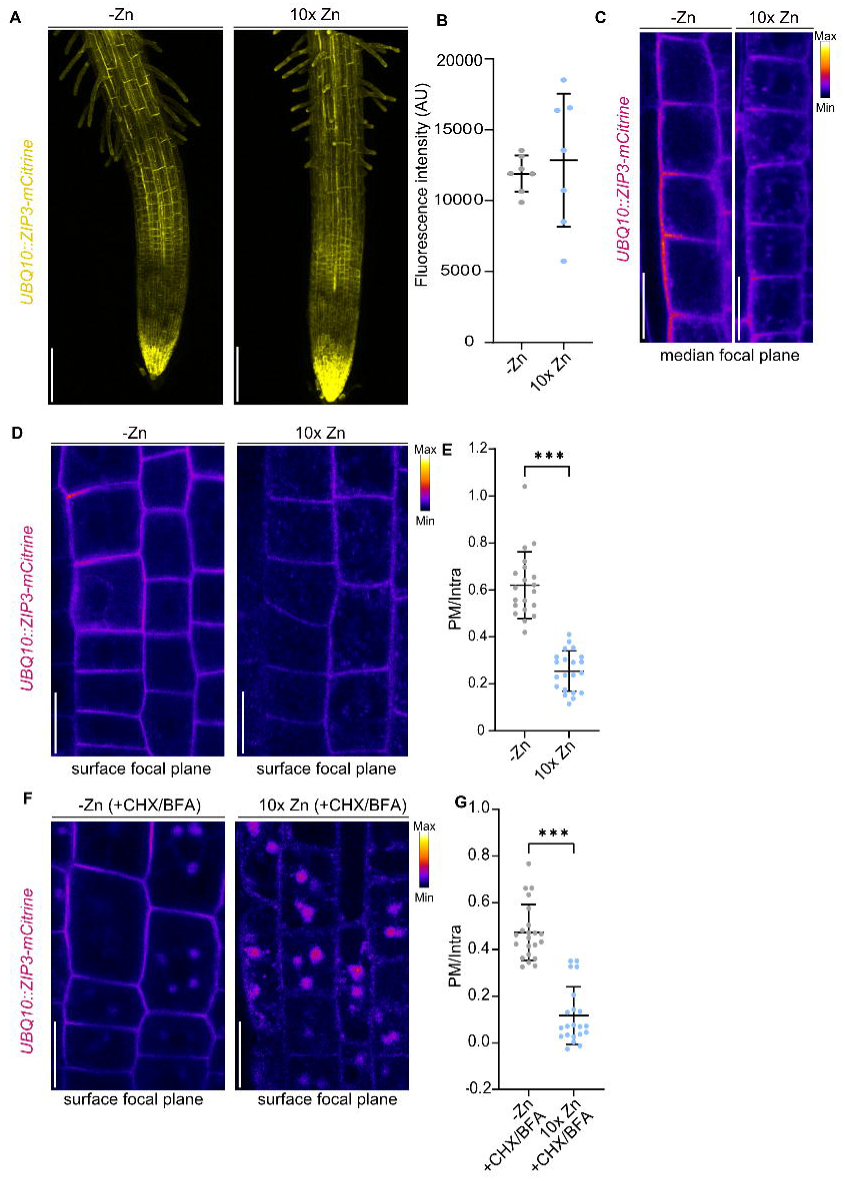
ZIP3 localization depends on Zn availability. **(A-G)** *UBQ10::ZIP3-mCitrine* seedlings were grown on agar plates containing Zn (15 µM) for 5 days, then transferred to liquid medium for 16 h under either Zn-deficient conditions (-Zn) or with a 10-fold excess of Zn (10x Zn; 150 µM) prior to imaging. **(A)** Representative maximum projections of 5-day-old roots expressing *UBQ10::ZIP3-mCitrine*. The mCitrine signal is shown in yellow. Scale bars: 125 µm. **(B)** Quantification of ZIP3-mCitrine signal intensity from maximum projection of several independent roots, as shown in panel A (n≥5). Data are represented as mean ±SD. No significant difference between conditions was observed using Mann-Whitney test (*P* < 0.05). **(C, D)** Close-up views of the elongation zone in roots shown in panel **A**, with images captured at the median focal plane of the epidermis **(C)** and at the surface of the epidermis **(D)**. The mCitrine signal is displayed using the Fire (LUT) color scale. Scale bars: 12.5 µm. **(E)** Quantification of the ratio of plasma membrane (PM) to intracellular (intra) of ZIP3-mCitrine signal (n≥20). Data are represented as mean ± SD, and statistical differences between conditions were determined using Mann-Whitney test (\**P* < 0.05; \*\**P* < 0.01; \*\*\**P* < 0.0001). **(F)** Surface views of epidermal cells from plants grown as described above and subsequently exposed to 100 µM CHX and 50 µM BFA prior to imaging. The mCitrine signal is displayed using the Fire (LUT) color scale. Scale bars: 25 µm. **(G)** Quantification of the ratio of PM to intracellular (intra) of ZIP3-mCitrine signal (n≥20) in plants treated as described in panel F. Data are represented as mean ± SD, and statistical differences between conditions were determined using Mann-Whitney test (\**P* < 0.05; \*\**P* < 0.01; \*\*\**P* < 0.0001).

## Discussion

Understanding the molecular mechanisms controlling metal acquisition by roots is essential for improving plant nutritional quality and growth, particularly under metal-limiting conditions. This is especially relevant for Zn, as Zn uptake mechanisms in dicotyledonous plants, in contrast to monocotyledonous, have been largely overlooked. To date, no Zn acquisition system in dicots has been unequivocally identified, despite the fact that dicot crops account for one-third of human plant-based nutrition (FAO, 2019). Given the widespread prevalence of Zn deficiency in human populations, enhancing Zn content and availability in crops is of paramount importance. Achieving this, however, requires the identification of key nutrient transporters, a detailed understanding of their substrate nature and specificity, their localization and regulation, as well as insights into how nutrient deficiencies impact the plant’s overall ionomic profile and the relationships between different elements. In this study, we sought to unravel the molecular mechanisms that enable Zn acquisition by the roots of dicot plants. To this end, we focused on the Arabidopsis ZIP family of transporters, aiming to identify those contributing to Zn acquisition from the soil. While many ZIP proteins are well-characterized for their role in metals transport in animals, bacteria and yeast, the ZIP family remains poorly understood in dicotyledonous plants, particularly in the context of metal uptake from the soil with the exception of IRT1 (Hantke, 2005; Lehtovirta-Morley et al., 2017; Ma & Gong, 1234; Roberts et al., 2021; Tuschl et al., 2016; Zhao & Eide, 1996a, 1996b). To bridge this knowledge gap, we first investigated the expression and promoter activity of *AtZIP* genes in Arabidopsis roots. By employing a highly fluorescent, non-mobile fluorophore, we were able to precisely map *ZIP* promoter activities to specific root cell types, providing insights into their potential functions. Through this approach, we successfully uncovered novel roles for 4 ZIP transporters involved in metal acquisition at the root periphery.

Firstly, we identified ZIP2 as a Cu transporter, and potentially a Mn transporter, involved in metal acquisition from the soil. While previous studies based on expression analysis and yeast complementation suggested ZIP2 could transport Cu in roots, *in vivo* evidence was still lacking (Wintz et al., 2003). Our physiological, confocal and ionomic analyses provide evidence that ZIP2 contributes to Cu acquisition in plants. Its vascular localization hints at a broader role beyond soil Cu acquisition, possibly in Cu partitioning between roots and shoots, similar to the dual role of IRT1 in Fe acquisition and distribution. Alongside ZIP2, other Cu transporters such as COPT1 and COPT2 are known to contribute to Cu^+^ acquisition in Arabidopsis, although their precise localization in root cell types remains unclear (Sancenón et al., 2004; Tiwari et al., 2017). The SPL7-(SQUAMOSA PROMOTER BINDING PROTEIN-LIKE7) dependent regulation of ZIP2, COPT1 and COPT2 underscores their significance in Cu acquisition, as the transcription factor SPL7 is a key regulator of Cu homeostasis (Bernal et al., 2012). Before Cu is taken up by root transporters, Cu^2+^ is reduced to Cu^+^ at the root surface by ferric reductases FRO3, FRO4 and FRO5, analogous to FRO2-mediated Fe^3+^ reduction (Bernal et al., 2012; Jeong & Connolly, 2009; Mukherjee et al., 2006). The promoter activities of *FRO3* in the vasculature and root tip, as well as FRO4 and FRO5 accumulation in the root tip, suggest that this zone is a hotspot for Cu uptake (Bernal et al., 2012; Mukherjee et al., 2006). COPT1 promoter activity in this region reinforces this idea. It is intriguing therefore, that ZIP2 does not accumulate in the root tip under our experimental conditions. This might indicate functional differences between ZIP2 and COPT1/COPT2 in Cu acquisition, as ZIP and COPT transporters facilitate the transport of Cu^2+^ and of Cu^+^ respectively (Peñarrubia et al., 2015). Whether ZIP2 interacts with FRO3/FRO4/FRO5 in specific tissues, similarly to the IRT1/FRO2 interaction remain an open question (Martín-Barranco et al., 2020). ZIP2 has also been proposed to mediate Zn transport in plants, based on ^65^Zn influx measurements in yeast and its ability to complement the *zrt1zrt2* yeast strain (Grotz et al., 1998; Milner et al., 2013). However, the absence of a Zinc Deficiency Response Element (ZDRE) in the *ZIP2* promoter, unlike those of *ZIP3* and *ZIP5*, suggests that Zn acquisition is not ZIP2’s primary function. ZDRE is known to be a key regulatory element targeted by the bZIP19/23 transcription factors, which are essential for Zn homeostasis (Assunção et al., 2010). Considering the broad specificity of ZIP transporters, it is not surprising that Mn over accumulated in the roots of *ZIP2::ZIP2-mCitrine* lines. Previous studies based on promoter-reporter lines and Mn quantification have already proposed that ZIP2 functions in Mn transport, particularly for Mn translocation from the root vasculature to the shoot (Milner et al., 2013). Our ionomic data further suggest that ZIP2 contribute to Mn acquisition from the soil, expanding the number of known Mn transporters in plants (Cailliatte et al., 2010). ZIP2 stands out from other ZIP family members because it is highly expressed and accumulates in the presence of metals, while its expression is repressed under Zn, Mn or Fe deficiencies conditions that typically induce other ZIP transporters (Figure 2B, Figure S1B,C, Milner et al., 2013; Wintz et al., 2003). This suggests that ZIP2 may play a role in basal metal uptake, functioning with lower affinity under deficient conditions, similar to the dual affinity nitrate transporters NRT1.1 and NRT2.1 (Y. Y. Wang et al., 2012). However, the low apparent *K_m_* for ZIP2 mediated Zn uptake does not fully support this hypothesis (Grotz et al., 1998). Surprisingly, ZIP2-mediated Zn accumulation, but not ZIP3-mediated uptake, is highly pH-dependent, raising questions about the conservation of Zn transport mechanisms among ZIP family members (Grotz et al., 1998). Thus, ZIP2’s function requires further investigation.

Secondly, we identified ZIP8 as an Fe transporter that localizes in a polar manner to the outer epidermis under Fe deficiency. Initially, *ZIP8* was proposed to be a pseudogene due to its inability to complement metal uptake mutants in yeast (*zrt1/zrt2Δ, fet3/fet4Δ, smf1Δ, ctr1/ctr3Δ*) (Milner et al., 2013). However, several studies have shown that *ZIP8* is upregulated in response to Fe deficiency and is part of the FIT-(FER-LIKE FE DEFICIENCY-INDUCED TRANSCRIPTION FACTOR) induced gene network, suggesting a role in Fe homeostasis (Bauer et al., 2004; Mai et al., 2016; Van De Mortel et al., 2006). Given its phylogenetic proximity to IRT1 and is specific induction under Fe deficiency, we investigated ZIP8’s role in Fe acquisition. Our data show that Fe accumulation is reduced in the roots of the *zip8* mutant, which strongly indicates that ZIP8 plays a direct role in Fe acquisition from the soil. The polar localization of ZIP8 to the outer epidermis reinforces this function. In contrast to IRT1, ZIP8-mCitrine signal is absent in the stele, indicating that ZIP8 does not participate in root-to-shoot partitioning. This suggests that ZIP8 and IRT1 have distinct, non-overlapping roles in Fe homeostasis. The absence of a macroscopic phenotype in *zip8* mutants and the lack of ZIP8 accumulation in the elongation zone, where IRT1 typically localizes, further support the notion that ZIP8 and IRT1 are not fully redundant. Notably, while *irt1* mutants accumulate high level of Fe in roots but exhibit Fe deficient in shoots, *zip8* mutants only show impaired Fe accumulation in roots, suggesting different physiological roles. Further investigation is needed to determine ZIP8’s substrate affinities and its relative contribution to total root Fe uptake in comparison to IRT1. It remains intriguing that overexpression of *ZIP8* or *IRT1* does not result in elevated Fe accumulation in roots. This could imply that endogenous levels of these transporters are already sufficient for Fe uptake, and increasing their expression or expanding their localization to other root tissues does not enhance Fe acquisition. Additionally, like IRT1, ZIP8 may have different metal affinities, which could explain the accumulation of non-Fe metals in overexpression lines.

Third, our findings provide multi-level evidence that in Arabidopsis, ZIP3, and to a lesser extent ZIP5, are key transporters involved in Zn acquisition in the outer root domain, similar to OsZIP9 and OsZIP5 in rice (Tan et al., 2020). Both ZIP3 and ZIP5 are strongly upregulated under Zn deficiency and localize partly to the PM of the outer root domain under these conditions. Notably, ZIP3 exhibits polar accumulation at the PM of the epidermis and the cortex, underscoring its central role in Zn uptake. Consistently, *zip3* mutants show reduced Zn accumulation compared to WT, while overexpression of both *ZIP3* and *ZIP5* enhances Zn accumulation in roots. Unlike IRT1, which when overexpressed leads to the accumulation of various metals (Barberon et al., 2011), ZIP3 and ZIP5 overexpression primarily increases Zn content, suggesting a higher specificity for Zn transport. This preference was supported by previous studies showing that Zn is the most effective competitor for ZIP3-mediated ^65^Zn uptake (Eide 1999). Given the similarity between ZIP3 and ZIP5, it is reasonable to hypothesize that ZIP5 also preferentially transport Zn. The localization of ZIP3 in the vasculature suggests that, in addition to its role in Zn acquisition, ZIP3 may also contribute to Zn partitioning between roots and shoots, similar to the role of IRT1 in Fe transport (Quintana et al., 2022). Although we did not observe a reduction in Zn translocation in the *zip3-1zip5-1* double mutant, this may be due to functional redundancy among other ZIPs, as several *ZIP* family members show promoter activity in the stele (Figure 2C, Lee et al., 2021; Milner et al., 2013). The role of the apoplast in metals homeostasis, particularly for Zn, is emerging as an important factor for regulating plants metal uptake (Peng et al., 2021; Zhong et al., 2024). A recent study has highlighted the significance of pectin methylesterification in the cell wall for Zn homeostasis (Zhong et al., 2024). The presence of ZIP3 and ZIP5 in the mature root cortex suggests that Zn uptake from the apoplast at the cortical level plays an essential role in Zn acquisition. However, the restriction of ZIP3 and ZIP5 to the epidermis in fully elongated roots may reflect variations in apoplastic metal accumulation along the root maturation gradient. While Zn transporters for acquisition in the outer root domain and for Zn efflux into the xylem have been identified, the radial transport of Zn between these two regions remains unclear. Two plausible, non-exclusive mechanisms could explain this process. First, based on promoter activity data, IRT3, ZIP1, ZIP7, ZIP10 and ZIP11 might facilitate Zn uptake at the endodermal level, suggesting that Zn diffuses through the apoplast to reach the endodermis or is exported into the apoplast from cortical cells by an efflux transporter. The presence of the Zn efflux transporter PCR2 in the cortex supports this hypothesis (Song et al., 2010). Alternatively, Zn may be taken up into the cytoplasm at the outer epidermis and transported radially via the ER localized transporter MTP2, moving cell-to-cell through plasmodesmata (PD), as already proposed (Sinclair et al., 2018). Although Zn transport though PD has yet to be demonstrated, this model would suggest that radial Zn transport does not depend on endodermal Zn transporters or polar efflux Zn transporter across root cell types, as proposed for the coupled transcellular pathway (Barberon & Geldner, 2014). The lack of identified polar Zn efflux carriers may indicate that symplastic transport is the preferred route for Zn radial movement.

In this study, we revealed that the transporters ZIP3 and ZIP8 localize in a polar fashion at the outer membrane of the epidermis. This observation suggests that polarity in nutrient transporters may be more widespread in plants than previously recognized, particularly in dicots, where only a few polar carriers have been identified (Robe & Barberon, 2023). This raises important questions regarding the polarity of others ZIP family member and nutrient carriers in general. Polarity has already been shown to be critical for the function of several carriers, and is believed to enable directional nutrient flow in roots (Barberon et al., 2014; Konishi et al., 2023; Yoshinari et al., 2019) as it is well established for PIN-(PIN FORMED-) mediated polar auxin transport(Feraru & Friml, 2008). The significance of ZIP3 and ZIP8 polarity in Zn and Fe acquisition, however, remains to be explored. It is plausible that their efficiency in metal acquisition is highly dependent on this precise polar localization. Whether ZIP3 and ZIP8 are regulated by mechanisms to those governing IRT1, such as FYVE1-(FYVE domain protein 1 also known as FREE1) mediated recycling and polarization in the outer epidermis, also warrants further investigation (Barberon et al., 2014). Zn, Fe, Cu and Mn often share common transporters and can influence each other’s uptake by competition, and their availability is often affected by similar soil conditions, such as high pH. Additionally, imbalances in one metal often impact the acquisition of others. In consequence, the localization and regulation of nutrient transporters must be tightly controlled. In our study, we observed that ZIP3 undergoes endocytosed in response to Zn excess, likely to prevent over-accumulation of Zn in cells, similar to IRT1 in Arabidopsis (Dubeaux et al., 2018). However, since ZIP3 lacks a histidine-rich stretch in its cytoplasmic domain – a key feature involved in Zn sensing for other transporters – the mechanism by which Zn levels are sensed remains unclear. It is possible that the single histidine between transmembrane domains (TD) 3 and 4 could play a role in this process. Interestingly, ZIP5 does contain a histidine-rich stretch between TD3 and 4, suggesting potential Zn binding and regulation in this region, hinting at similar regulatory mechanisms.

Extending from knowledge of ZIP transporters in animals, the polarity of nutrient carriers is not unique to plants. In animal, polarized localization of ZIP family transporters is known to be crucial for maintaining metal homeostasis in specific tissues (Liuzzi & Cousins, 2004; Taylor & Nicholson, 2003). For example, hZIP4 in humans is polarized at the apical membrane of intestinal cells, facilitating directional Zn uptake from the diet (Wang et al., 2004). This polarization is tightly regulated and influenced by Zn availability, similar to what we observed for ZIP3 in Arabidopsis. Furthermore, substrate-induced endocytosis is a well-documented mechanism for controlling transporter activity in both plants and animals. In mammals, hZIP4 undergoes Zn-induced endocytosis to reduce Zn uptake when cellular levels are sufficient (Zhang et al., 2020), a process similar to the endocytosis observed for ZIP3 under Zn excess. The presence or absence of specific histidine motifs in these transporters often governs their sensitivity to metal levels and their endocytic regulation. Such insights from animal systems highlight the potential for conserved mechanisms across kingdoms, emphasizing the importance of polarity and substrate-induced endocytosis in maintaining nutrient homeostasis in both plants and animals.

In conclusion, by investigating the spatial dynamics of ZIP transporters, we highlight the need to further explore transporter localization as a critical factor for understanding their function and the regulatory mechanisms that optimize nutrient acquisition and homeostasis in plants.

## Material and methods

### Plant Material

All experiments were conducted using *Arabidopsis thaliana* wild type (WT) Columbia-0 background. Mutants and transgenic plants used in this study are as follow: *zip2-1* (SALK_094937, Milner et al., 2013), *zip3-1* (salk_35_B08, Nishida et al., 2015), *zip5-1* (SALK_009007C, Wu et al., 2009), all of which were obtained from the Nottingham Arabidopsis Stock Center (NASC). *irt1-2* was obtained from (Varotto et al., 2002). Additionally, the following mutants were generated for this study using CRISPR-Cas9 technology: *zip2-2_cr_, zip3-2_cr_, zip5-2_cr_, zip8-1_cr_* and *zip8-2_cr_.* Transgenic lines generated for this research include: *pIRT1::NLS-3xmVENUS, pIRT2::NLS-3xmVENUS, pIRT3::NLS-3xmVENUS, pZIP1::NLS-3xmVENUS, pZIP2::NLS-3xmVENUS, pZIP3::NLS-3xmVENUS, pZIP4::NLS-3xmVENUS, pZIP5::NLS-3xmVENUS, pZIP6::NLS-3xmVENUS, pZIP7::NLS-3xmVENUS, pZIP8::NLS-3xmVENUS, pZIP9::NLS-3xmVENUS, pZIP10::NLS-3xmVENUS, pZIP11::NLS-3xmVENUS, pZIP12::NLS-3xmVENUS, pZIP2::ZIP2-mCitrine, pZIP3::ZIP3-mCitrine, pZIP5::ZIP5-mCitrine, pZIP8::ZIP8-mCitrine, UBQ10::ZIP2-mCitrine, UBQ10::ZIP3-mCitrine, UBQ10::ZIP5-mCitrine, UBQ10::ZIP8-mCitrine, pZIP2::IRT1, pZIP3::IRT1, pZIP5::IRT1, p35S::ZIP3, p35S::ZIP5* and *p35S::IRT1.* The construct *pIRT1* was ordered from NASC (N2106308).

### Constructs

Plasmids were generated using Gibson and Multisite Gateway cloning techniques (Thermo Fisher Scientific). A list of primers utilized for cloning is provided in Table S2. *ZIP* promoter sequences upstream of the ATG start codon were amplified from Arabidopsis Col-0 genomic DNA and inserted into a modified *pDONRP4-P1R* (Thermo Fisher Scientific) via Gibson assembly. Briefly, the *ccdB* and the *CmR* cassettes between the AttP1R and attP4 recombination sites were removed in *pDONRP4-P1R* and replaced by EcoRV, BglII, XbaI, BamhI cloning sites. For the generation of promoter-reporter constructs (*promoter::NLS-3xmVenus*), entry plasmids containing the promoter regions were recombined with *pDONRL1-NLS-3xmVenus-L2* and *pEN-R2-tHSP18.2-L3* into the destination vector *pFR7m34GW* (Kalmbach et al., 2023). To study endogenous protein localization, *ZIP* genomic sequences were amplified from Col-0 genomic DNA and inserted by Gibson in a modified *pDONR221,* containing cloning sites instead of *ccdB* and *CmR* cassettes. The mCitrine fluorescent protein was inserted into the second extracellular loop of each ZIP transporter using Gibson assembly. For ZIP2-mCitrine, mCitrine was inserted 153 bp downstream of the ATG start codon; for ZIP3-mCitrine, it was inserted 95 bp from ATG; for ZIP5-mCirinet, 89 bp from ATG; and for ZIP8-mCitrine, 87 bp from ATG. The final destination vectors for expression in plants were obtained using multisite Gateway recombination system (Life Technologies), employing the *pFR7m34GW* destination vector along with the various *pDONR221-ZIP-mCitrine* entry clones to generate the *ZIP::ZIP-mCitrine* constructs. For promoter swap analysis, entry vector containing the promoters of *ZIP2*, *ZIP3* and *ZIP5* were recombined with *pDONRL1-gIRT1-L2* and *pEN-R2-tHSP18.2-L3* into the destination vector *pFR7m34GW.* Vectors carrying *ZIP2/3/5::IRT1* were transformed in *zip3-1zip5-1* background to evaluate IRT1 function in Zn uptake. For CRISPR/Cas9 mutant generation, single guide RNA (sgRNA) for Cas9 were designed using webtool Benchling (https://www.benchling.com). Pairs of annealed oligonucleotides of the sgRNA were cloned into the BbsI linearized entry vector recombined into the destination vector containing Cas9 expression cassette controlled by the *PcUBi4-2* promoter and a FastRed selection marker (Ursache et al., 2021). To generate the *zip2-2_cr_* mutant, a single sgRNA targeting the 5’ genomic region was combined with the *ZCas9i* controlled by the *pEC1.2* promoter (Grützner et al., 2021). For the generation of the *zip3-2_cr_*, *zip5-2_cr_*, *zip8-1_cr_* and *zip8-2_cr_*, triple sgRNA targeting the 5’ genomic region were employed with *Cas9*. All constructs were transformed into Agrobacterium strain GV3101 by electroporation and subsequently used for Arabidopsis plant transformation via the floral-dip method. T1 plants were first selected based on the red fluorescence of transformed seeds and fluorescent mono-insertional T2 were subsequently selected. All experiments were performed on mono-insertional T2 after red fluorescence sorting unless otherwise specified in figure legends. CRISPR-induced mutation experiments were performed on T3 or T4 homozygous lines.

### Growth conditions

For *in vitro* culture, seeds were surface sterilized in 70 % ethanol for 5 minutes with agitation, followed by 3 rinses with absolute ethanol before drying. The sterilized seeds were then sown on square plates containing half-strength Murashige and Skoog (MS) based media with 0.8% agar (Duchefa) and no sucrose. For metal deficiency or sufficiency, plants were grown in the presence (+Zn, +Cu, +Fe +Mn) or absence (-Zn, –Cu,-Fe, –Mn) of 15 µM Zn, 0.05 µM Cu, 50 µM Fe and 50 µM Mn provided as ZnSO_4_, CuSO_4_, Fe-EDTA and MnSO_4_ respectively. Additional nutrients were supplied at following concentration: 9.4 mM KNO_3_, 10.3 mM NH₄NO₃, 625 mM KH_2_PO_4_, 1.49 mM CaCl_2_, 0.75 mM MgSO_4_, 50 µM H_3_BO_3_, 0.5 µM Na_2_MoO_4_, 2.5 µM KI, 0.05 µM CoCl_2_. After stratification in the dark for 3 days at 4°C, the plates were transferred to growth chamber under continuous day conditions (light intensity ∼100 μE) at 22 °C with ∼50% humidity and the plants grown vertically. For physiological analysis, root length, FW and chlorophyll contents were assessed in 10-day-old plants grown on control or metal-deficient agar plates. For elemental analysis, plants were grown either on agar control media for three weeks or in soil for the same duration before harvesting. For mRNA relative quantification, 7-days old seedlings were directly sown and grown on control or metal deficient plates before harvesting. Live-microscopy analyses were conducted on 5-day-old seedlings. For promoter activity, expression analysis and protein localization, plants were directly sown and grown for 5 days on control or metal-deficient agar plate. For short-term metal deficiency or excess treatments, 5-day-old plants were transferred to liquid medium without Zn (-Zn) or with an excess of Zn (10xZn; 150 µM of Zn provided as ZnSO_4_) for 16 hours before imaging. CHX was applied to plants at a final concentration of 100 μM for 1 hour before treatment with 50 μM BFA, which was maintained for the duration of the BFA treatments (3 hours). For amplification and experiments in soil, plants were grown in long-day conditions (16 hours of light, 8 hours of darkness) with a light intensity of 150-180 μE at 60-70% humidity, and at 20 ± 2 °C.

### Confocal Microscopy and image analysis

Cell walls were stained with a 10 μg.ml^-1^ solution of propidium iodide (PI; Sigma-Aldrich, Cat# P4170) for 10 minutes, while the PM and endocytic compartments were stained with a 4 µM solution of the styryl dye FM4-64 (ThermoFischer scientific, Cat# T13320) for 10 minutes. Zn visualization was conducted with 10 µM Zinpyr-1 (Abcam, Cat# 145349) staining as described by Sinclair et al., 2007. To prevent excessive Zn accumulation in the roots and minimize fluorescent signal saturation, plants were grown under control conditions with 1 µM Zinc (+Zn) instead of 15 µM.

Confocal laser scanning experiments were conducted using a Zeiss LSM 780 system. The excitation (ex) and detection (em) settings were configured as follows: mVenus/mCitrine ex: 514 nm, em: 515 – 570 nm; propidium iodide (PI) ex: 514 nm, em: 586 – 679 nm; Zinpyr-1: ex: 488nm, em: 495 – 535 nm; and FM4-64 ex: 514 nm, em: 650 – 742 nm. Images were captured using a 40X objective for Z-stack imaging and a 20X dry objective for longitudinal views. All images were processed using the Fiji software (LUT, orthogonal views, maximum projections, merge, fluorescence quantification etc) (Schindelin et al., 2012). For Z-stack images of promoter-reporter lines, protein localization and Zinpyr-1 staining, the Z-stack images were resliced vertically to generate transversal views. For promoter reporter lines only, a maximum projection of the mVenus marker channel was then merged (3D; X, Y, Z) with a representative single stack of the PI-stained cell wall channel (2D; X, Y). For T1 lines, at least five independent lines were observed and for T2 lines, at least two independent mono-insertional lines were observed and a representative line is shown in the manuscript. For Figure 7, PM/intra signal were calculated as described in Spielmann et al., 2023. The fluorescence intensity of Zinpyr-1 in the endodermis was determined using identical regions of interest (ROI). The Zinpyr-1 signal in several endodermal cells from several independent roots was measured to obtain the mean Zinpyr-1 fluorescence intensity.

### qRT-PCR

For gene expression analysis, roots from approximately 40 7-day-old seedlings were harvested and pooled to form one biological replicate. RNA were extracted using a TRIzol-adapted RNeasy MinElute CleanupKit (Qiagen). Subsequently, RNA were reverse transcribed with a Thermo Scientific Maxima First Strand cDNA Synthesis Kit following the manufacturer’s protocol. Quantitative Real-time PCR was conducted on an Applied Biosystems QuantStudio5 thermo-cycler using Applied Biosystems SYBR Green master mix. The relative expression of *ZIP* genes was quantified with the 2^−ΔΔCt^ method with *Clathrin* as a reference gene (AT5G46630). A list of primers used for qRT-PCR is provided in table S2. Each experiment included 3 technical and at least three biological replicates.

### Ionomic analysis

Three week-old seedlings were used to quantify metal content. Shoots and roots were harvested separately and desorbed by washing in 10 mM Na_2_EDTA for 10 minutes, followed by 3 rinses with ultrapure Milli-Q water for 1 minutes each. After desorbing, the tissues were blotted dry with paper towels and placed at 65°C overnight. Once dried, the tissues were ground, and 5-10 mg of dry tissue per biological replicate was mixed with 750 µl of nitric acid (65% [v/v]) and 250 µl of hydrogen peroxide (30% [v/v]). The samples were left at room temperature overnight and then mineralized at 85°C for 24 hours. Following mineralization, 4 ml of Milli-Q water was added to each sample. Metal content was measured using Inductively Coupled Plasma Optical Emission Spectrometry (ICP-OES). Four to five biological replicates were analyzed for each genotype.

### Chlorophyll quantification and F_v_/F_m_ measurements

Chlorophylls were extracted from 3 weeks (only for figure 1C) or 10-day-old plants tissues by incubating sample overnight at 4°C in 10 volumes of dimethylformamide (DMF) (v/w). Following extraction, samples were centrifugated, and chlorophyll content was quantified using a NanoDrop spectrophotometer at wavelength 647 nm and 664 nm. Chlorophyll concentrations were calculated according to the methods previously described by Porra et al., 1989. For Fv/Fm measurement, plants were dark-adapted for 15 minutes before measurements were taken with a FluorCam 800 MF, as described in Leonardelli et al., 2024. The maximum quantum efficiency of photosystem II (F_v_/F_m_) was measured by applying saturating pulses of 2000 µmol m^−2^ s^−1^(white light, 400 to 720 nm) for 960 ms, followed by detection using orange-red light pulses (620 nm,10 µs). All measurements were performed at room temperature.

### Phylogenetic tree

Protein sequences for *Arabidopsis thaliana* ZIP family members were retrieved from TAIR (https://www.arabidopsis.org). Phylogenetic relationships were defined using Phylogeny.fr (http://www.phylogeny.fr/).

### AlphaFold and AlphaFold2 protein interaction prediction

Wild type ZIP and ZIP-mCitrine models were predicted using AlphaFold (a) (https://alphafold.ebi.ac.uk/) or the ColabFold v1.5.5: AlphaFold2 using MMseqs2 (b). (Jumper et al., 2021; Kim et al., 2024; Varadi et al., 2022). The number of recycles were set to 3 for ColabFold predictions.

### Statistical Analysis

All statistical analyses were conducted using GraphPad Prism software or within the R environment (R Core Team 2023). Data were first assessed for normality and homoscedasticity using the Shapiro-Wilk test and Brown-Forsythe test, respectively. For data meeting these assumptions, pairwise comparisons were performed using a two-tailed Student’s *t*-test, while multiple comparisons were conducted using a one-way ANOVA followed by a Tukey’s post hoc test. When the assumptions of normality or homoscedasticity were violated, non-parametric tests were employed. In such cases, pairwise comparisons were carried out using the two-tailed Mann-Whitney U test, and multiple comparisons were analyzed using the Kruskal–Wallis test followed by Dunn’s post hoc test or a non-parametric version of Tukey’s test (R). A significance threshold of *P* < 0.05 was applied in all tests. All experiments were repeated at least 3 independent times with the exception of experiments in Figure S4B, S6C and representative experiments are presented in the manuscript.

### Accession Numbers

The sequence data used in this article (for generating constructs and for qRT-PCR) can be found in the GenBank/EMBL databases under the following numbers: *IRT1* (*AT4G19690*), *IRT2* (*AT4G19680*), *IRT3* (*AT1G6096*0), *ZIP1* (*AT3G12750), ZIP2* (*AT5G59520*), *ZIP3* (*AT2G32270*), *ZIP4* (*AT1G10970*), *ZIP5* (*AT1G05300*), *ZIP6* (*AT2G30080*), *ZIP7* (*AT2G04032*), *ZIP8* (*AT5G45105*), *ZIP9* (*AT4G33020*), *ZIP10* (*AT1G31260*), *ZIP11* (*AT1G55910*), *ZIP12* (*AT5G62160*).

## Acknowledgments

We would like to thank Lothar Kalmbach, Satoshi Fujita, Robertas Ursache and Joop E.M. Vermeer for sharing plasmids We also thanks Léa Jacquier for performing statistics in the R environment and Emilie Demarsy for her assistance with chlorophyll quantification and fluorescence measurements as well as Isabelle Fleury for their help in plant propagation. We thank Sylvain Philippe Loubery for help with the microscopes and Anna Puzyrko, Elliot Bowles and Laura Pillard-Tapia for technical assistance during their internships. Special thanks to Sandrine Chay and Stephane Mari from the Multi-Elemental Analyses Service (SAME) at the Institute for Plant Sciences of Montpellier (IPSiM). We also extend our thanks to the Photonic Biomaging Center at the University of Geneva. This work was supported by the Swiss National Science Foundation (<u>PCEGP3_187007</u>) warded to MB and by the state of Geneva.

## Supplemental figures

**Figure S1.**
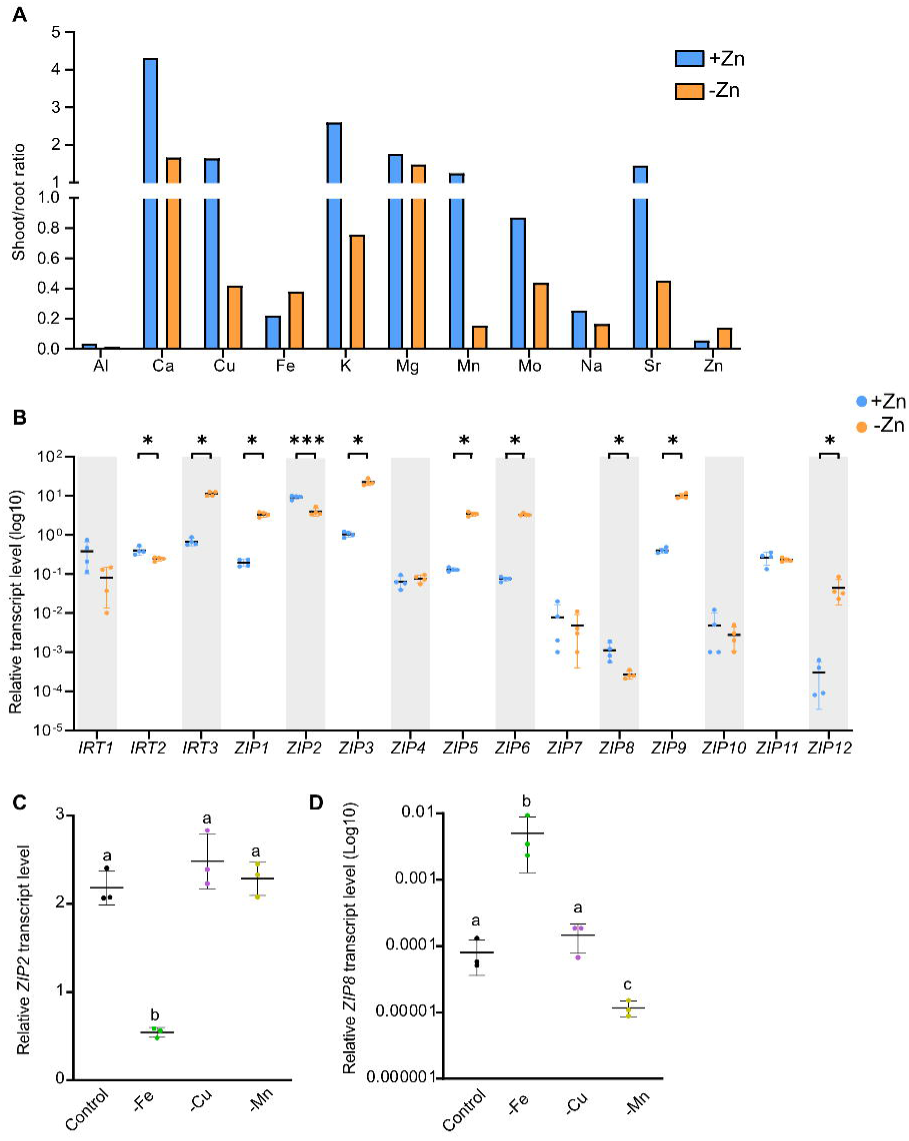
Zinc deficiency affects shoot-to-root elemental ratios and *ZIP* gene expression. **(A)** Shoot-to-root ratios of the average elemental content in WT plants grown for 3 weeks on agar plate in presence (+Zn) or absence (-Zn) of zinc. Values correspond to the ratios from 3 independent pools of at least 10 individual shoots and roots (See Table S1 for numerical values). **(B)** Relative transcript levels of the 15 *ZIP* members in WT roots from plants grown for 1 week under +Zn or –Zn conditions. Each dot represent one biological replicate. Data are represented as mean ± SD (n=4, each sample is a pool of at least 40 seedlings). Statistical differences were determined using Student’s *t*-test or Mann-Whitney test (\**P* < 0.05; \*\**P* < 0.01; \*\*\**P* < 0.0001). **(C-D)** Relative transcript levels of **(C)** *ZIP2* and **(D)** *ZIP8* in WT roots from plants grown under metal-sufficient (Control) or metal-deficient (-Fe, –Cu, –Mn) conditions. Data are represented as mean ± SD (n=3, each sample is a pool of at least 40 roots). Different letters indicate statistically significant differences between conditions determined by one way ANOVA followed by Tukey post hoc test or Kruskal–Wallis test followed by non-parametric Tukey test (*P* < 0.05)

**Figure S2.**
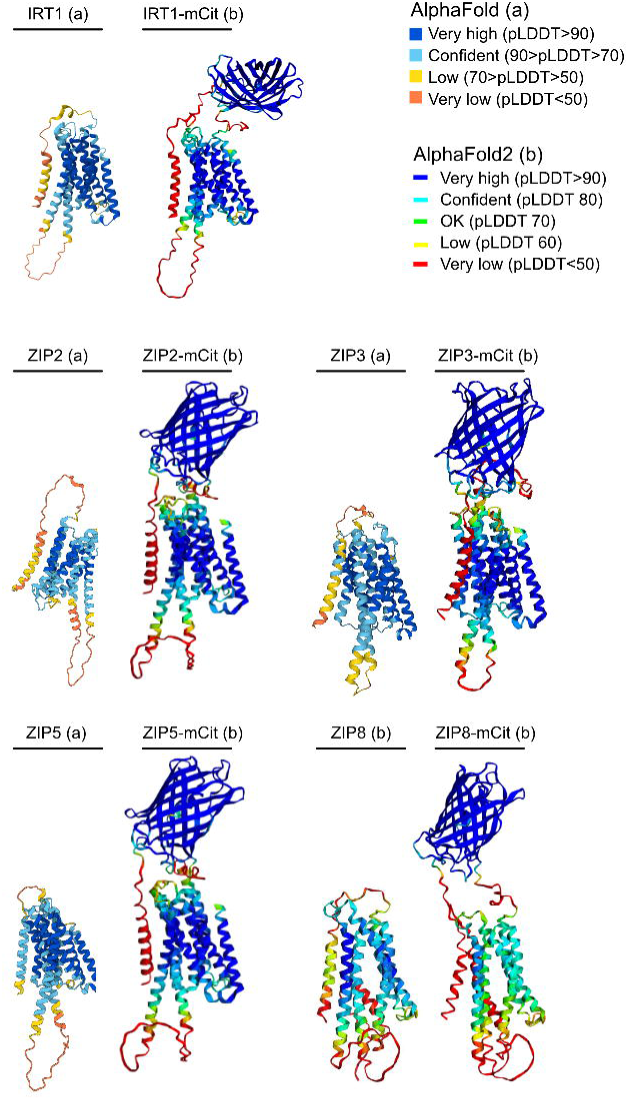
AlphaFold models. Predicted 3D structures of WT IRT1, ZIP2, ZIP3, ZIP5, ZIP8 and their fluorescent versions were generated using AlphaFold (a) or AlphaFold2 (b) respectively. For ZIP5 and ZIP8, we used *ZIP5.1* and *ZIP8.1* transcript versions for structure prediction.

**Figure S3.**
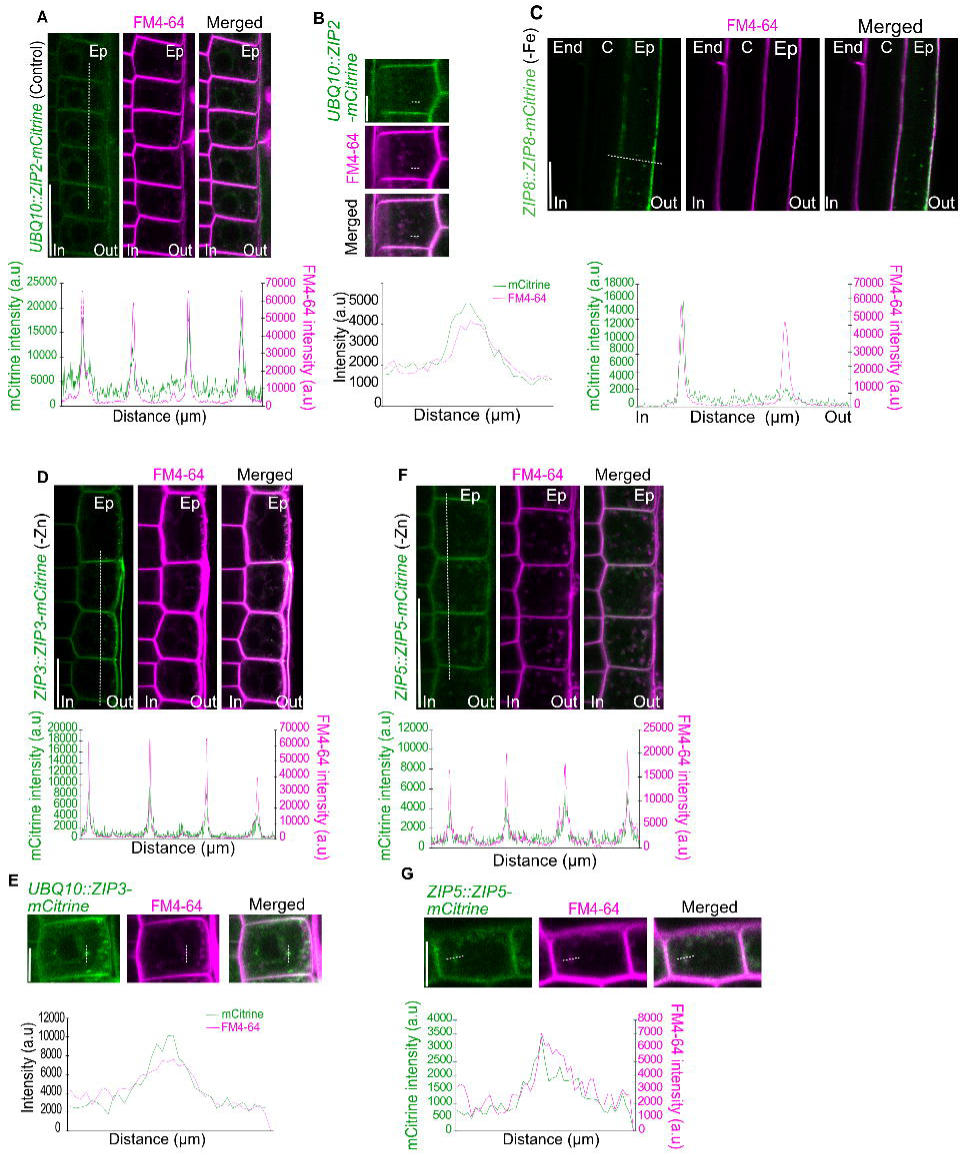
Co-localization of ZIP2, ZIP3, ZIP5 and ZIP8-mCitrine with FM4-64. **(A, C, D, F)** Co-localization of ZIP2, ZIP3, ZIP5 and ZIP8-mcitrine with FM4-64 in the epidermis of the root elongation zone (A, D, F) or the differentiated root (C), imaged 10 minutes after FM4-64 staining (top panels). The fluorescence signals of mCitrine and FM4-64 are plotted along the dashed lines in each image to show overlap (bottom panels). Scale bars: 25 µm. **(B, E, G)** Co-localization of ZIP2, ZIP3 and ZIP5-mCitrine with FM4-64 at endosomes in the epidermis of the elongation zone, 10 minutes after FM4-64 staining (top panels). The fluorescence intensity profiles of mCitrine and FM4-64 are plotted across the dashed lines in each image (bottom panels). Scale bars: 6.25 µm. a.u; arbitrary units.

**Figure S4.**
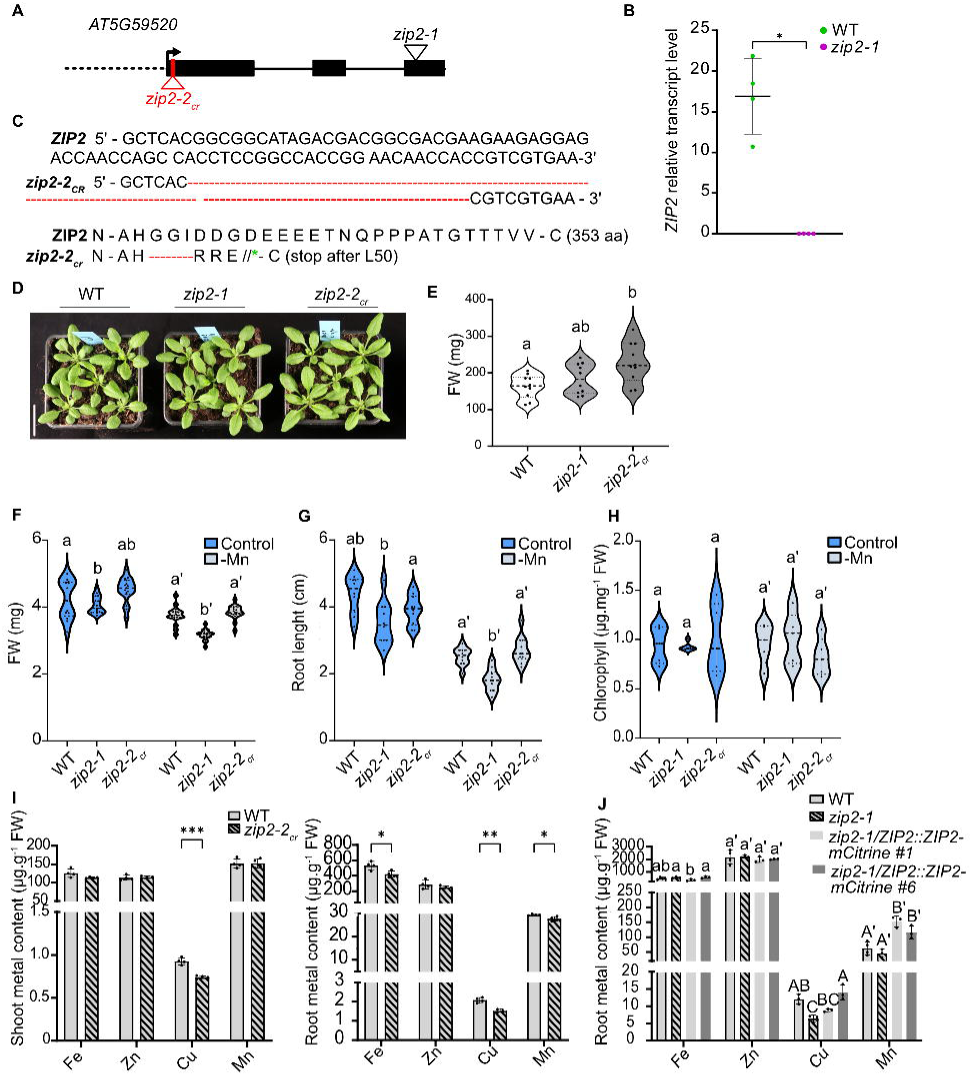
Characterization of *ZIP2* mutants and complementation lines. **(A)** Schematic representation of the genomic structure of *AtZIP2,* showing the locations of the T-DNA insertion in *zip2-1* and the CRISPR-induced mutation in *zip2-2_cr_*. Black boxes represent exons, the black lines represent introns, the dashed line indicates the promoter region, and the black angled arrow marks the START codon. **(B)** Relative *ZIP2* transcript levels in WT and *zip2-1* plants grown for 1 week on control agar plates is shown (n=4, each biological replicate consisting of a pool of at least 40 roots). Data are represented as mean ± SD, and statistical significance was determined using the Mann-Whitney test (\**P* < 0.05; \*\**P* < 0.01; \*\*\**P* < 0.0001). **(C)** Partial genomic (top) and protein (bottom) sequence of *ZIP2* in WT and the *zip2-2_cr_* mutant. The *zip2-2_cr_* allele results in a truncated protein due to a premature stop codon, producing only 50 amino acids (aa) compared to the 353 aa in the WT. **(D)** Phenotype of WT, *zip2-1* and *zip2-2_cr_* plants grown for 3 weeks in soil under long-day conditions. Scale bar: 2cm. **(E)** Quantification of fresh weight (FW) for plants grown as in panel C (n≥10). In the violin plots dashed lines represent the median and dotted lines represent the first and third quartiles. Different letters indicate statistically significant differences between different genotypes after one-way ANOVA followed by Tukey’s post hoc test (*P* < 0.05). (**F-H**) Violin plots representing **(F)** FW (n≥15), **(G)** root length (n≥15) and **(H)** chlorophyll (n≥5) content of WT, *zip2-1* and *zip2-2_cr_* plants grown for 10 days under control (+Mn) and manganese-deficient (-Mn) conditions. In all panels, dashed lines represent the median, and dotted lines mark the first and third quartiles. Different letters indicate statistically significant differences between different genotypes using one-way ANOVA followed by Tukey’s post hoc test or Kruskal– Wallis test followed by Dunn’s test where appropriate (*P* < 0.05). **(I)** Fe, Cu, Mn and Zn content in shoots (left) and roots (right) of WT and *zip2-2_cr_* mutant grown vertically for 3 weeks on control agar plates. Each replicate consisted of 3 to 4 plants pooled together (n=4). The data are presented as mean ± SE and significant differences between WT and *zip2-2_cr_* were determined using the Student’s *t*-test or Mann-Whitney test (\**P* < 0.05; \*\**P* < 0.01; \*\*\**P* < 0.0001). **(J)** Metal content (Fe, Cu, Mn, Zn) in roots of WT, *zip2-1*, and *zip2-1/ZIP2::ZIP2-mCitrine* complementation lines, grown 3 weeks on control agar plates. For each biological replicate, 3-4 plants were pooled (n=3-4). Different letters indicate statistically significant differences between genotypes using one-way ANOVA followed by Tukey’s post hoc test or Kruskal– Wallis test followed by Dunn’s test where appropriate (*P* < 0.05).

**Figure S5.**
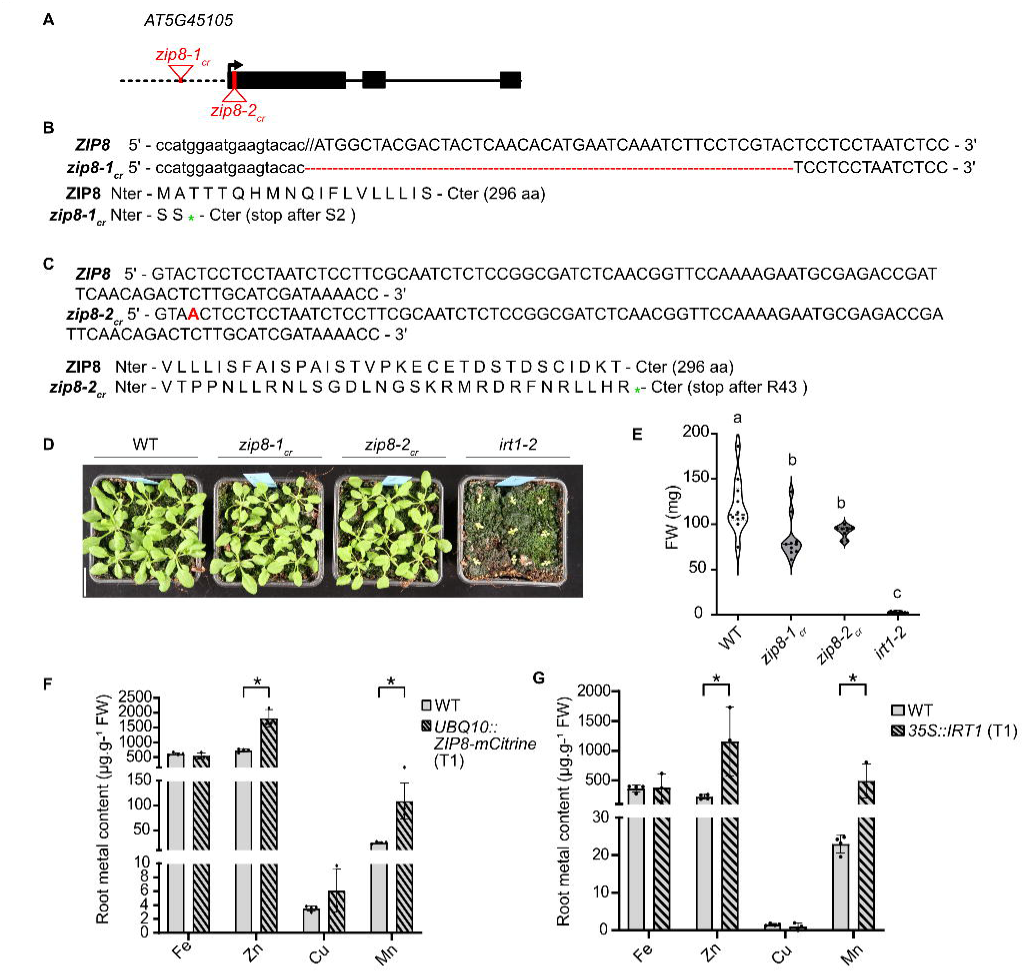
Characterization of *ZIP8* mutants and complementation lines. **(A)** Schematic representation of the genomic structure of *AtZIP8* showing the location of the *zip8-1_cr_* and *zip8-2_cr_* mutations. Black boxes represent exons, the black lines represent introns, the dashed line indicates the promoter region, and the black angled arrow marks the START codon. **(B, C)** Partial *ZIP8* genomic sequence (top) and protein sequence (bottom) for WT, *zip8-1_cr_ and zip8-2_cr_* mutants. WT *ZIP8* encodes a 296 aa protein, while *zip8-1_cr_* has a theoretical protein of 2 aa and*. zip8-2_cr_* encodes only 43 aa due to a premature stop codon. **(D)** Phenotype of WT, *zip8-1cr* and *zip8-2_cr_* mutants grown for 3 weeks on soil under long-day conditions. Scale bar: 2cm. **(E)** FW of plants grown as described in panel D. Different letters indicate statistically significant differences between different genotypes using Kruskal–Wallis test followed by non-parametric Tukey test (*P* < 0.05). **(F-G)** Metal content (Fe, Cu, Mn, Zn) in roots of **(F)** *UBQ10::ZIP8-mCitrine* lines and **(G)** *35S::IRT1* lines grown for 3 weeks on vertical control agar plates. Four independent T1 plants were pooled to form one biological replicate (n=3-4). Data are represented as mean ± SE, with statistical differences determined using Student’s *t*-test or Mann-Whitney test (\**P* < 0.05; \*\**P* < 0.01; \*\*\**P* < 0.0001).

**Figure S6.**
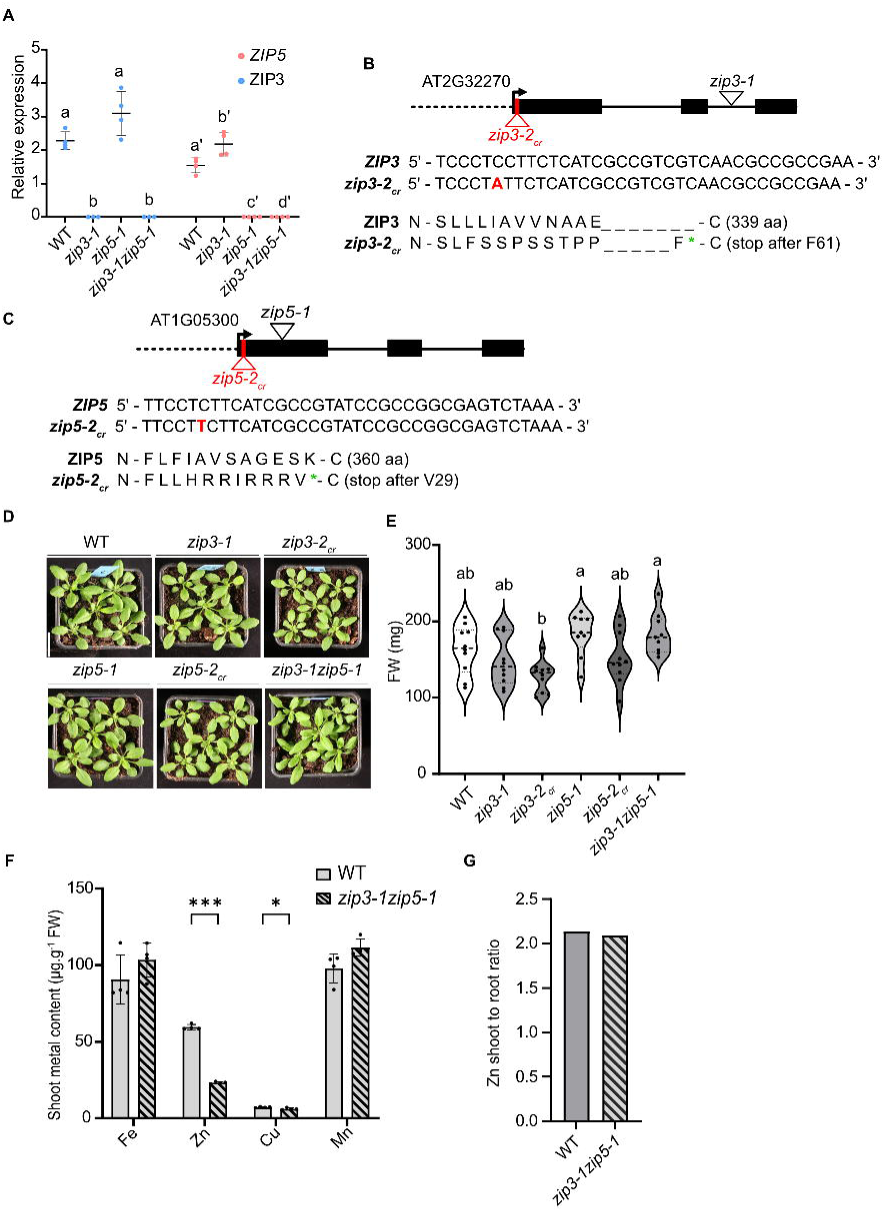
Genomic structure and phenotype of *zip3* and *zip5* mutants. **(A)** Relative *ZIP3* and *ZIP5* transcript levels in WT, *zip3-1*, *zip5-1* and the double mutant *zip3-1zip5-1* grown for 1 week under Zn-deficient conditions. Four biological replicates were analyzed, with each replicate consisting of a pool of at least 40 root seedlings. Data are represented as mean ± SD, with statistical differences between genotypes per individual gene determined using Kruskal–Wallis test followed by non-parametric Tukey’s test (*P* < 0.05). **(B, C)** Schematic representation of the genomic structure of *AtZIP3* and *AtZIP5,* showing the location of the T-DNA insertion in *zip3-1*, *zip5-1* and the *zip3-2_cr_ and zip5-2-2_cr_* CRISPR-induced mutations. Black boxes represent exons, the black lines represent introns, the dashed line indicates the promoter region, and the black angled arrow marks the START codon. Partial *ZIP3* and *ZIP5* genomic sequences (top) and protein sequences (bottom) for WT, *zip3-2_cr_* and *zip5-2_cr_* mutants. WT ZIP3 encodes a 339 aa protein, while *zip3-2_cr_* encodes a truncated protein of 61 aa. Similarly, WT ZIP5 encodes a 360 aa protein, while *zip5-2_cr_* produced a truncated protein of 29 aa **(D)** Representative phenotype of WT, *zip3-1, zip3-2cr, zip5-1, zip5-2cr and zip3-1zip5-1* plants grown on soil for 3 weeks under long-day conditions. Scale bar: 2 cm. **(E)** Violin plots showing the distribution of the FW of plants grown as described in panel D (n≥10). In the violin plots dashed lines represent the median and dotted lines represent the first and third quartiles. Different letters indicate statistically significant differences between genotypes using one-way ANOVA followed by Tukey’s post hoc test (*P* < 0.05). **(F)** Metal content (Fe, Cu, Mn, Zn) in shoots of WT and *zip3-1zip5-1* mutants grown for 3 weeks on soil under long-day conditions. Three to four plants were pooled to form one biological replicate (n=4). Data are represented as mean ± SE, with statistically significant differences between genotypes determined using Student’s t-test (\**P* < 0.05; \*\**P* < 0.01; \*\*\**P* < 0.0001). (**G**) Zn shoot/root ratio in WT and *zip3-1zip5-1* grown three weeks on soil under long day conditions.

**Figure S7.**
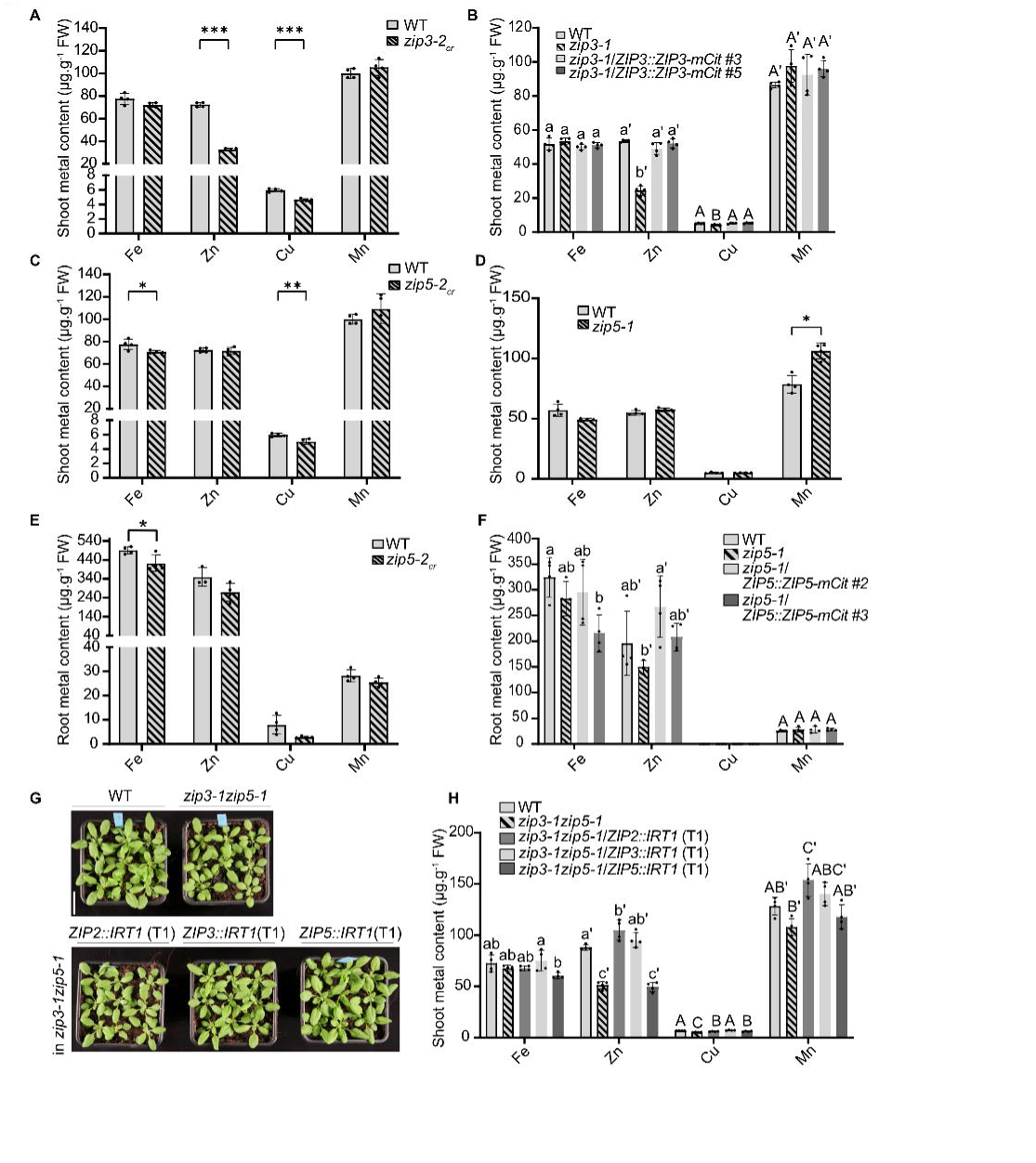
ZIP3 and ZIP5 individual contribution in metal acquisition. **(A-F)** Metal content (Fe, Cu, Mn, Zn) in shoots (A-D, H) and roots (E,F). (**A-F**) Three to four plants were pooled to form one replicate (n=4). (**A**, **C**, **D, E**) Statistical differences between genotypes for a given element were determined using Student’s *t*-test or Mann-Whitney test (\*\*\**P* < 0.0001; \*\**P* < 0.01; \**P* < 0.05). (**A-D, H**) Plants were grown 3 weeks on soil under long-day conditions. (**A**) Metal content in WT and *zip3-2_cr_* plants. (**B**) Metal content in WT, *zip3-1* and two complementation lines *zip3-1/ZIP3::ZIP3-mCitrine.* Different letters indicate statistically significant differences between genotypes for a given element using one-way ANOVA followed by Tukey’s post hoc test (*P* < 0.05). (**C**) Metal content in WT and *zip5-2_cr_* plants. (**D**) Metal content in WT and *zip5-1* plants. **(E)** Metal content in WT and *zip5-2_cr_* plants. **(F)** Metal content in WT, *zip5-1* and two complementation lines *zip5-1/ZIP5::ZIP5-mCitrine* grown for 3 weeks on agar plate. For the complementation lines, 3-4 independent T2 plants were pooled to form one biological replicate. Different letters indicate statistically significant differences between genotypes for a given element using one-way ANOVA followed by Tukey’s post hoc test (*P* < 0.05). **(G)** Phenotype of WT, *zip3-1zip5-1* and *zip3-1zip5-1/ZIP_2/3/5_::IRT1* lines. For each transgenic line, 9 independent T1 were grown per pot. **(H)** Metal content of the plants grown as described in panel G. For each biological replicate 3-4 independent T1 plants were pooled (n=4). Different letters indicate statistically significant differences between genotypes for a given element using one-way ANOVA followed by Tukey’s post hoc test (*P* < 0.05).

**Figure S8.**
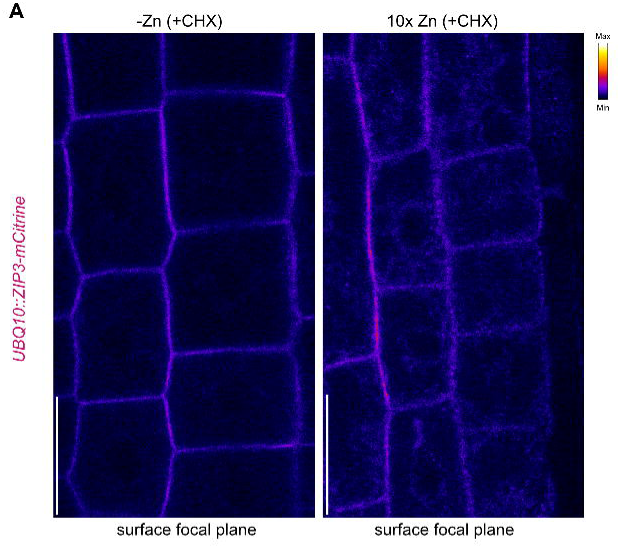
ZIP3 localization depends in Zn availability. Surface views of root epidermal cells from 5-day-old plants expressing *UBQ10::ZIP3-mCitrine* grown on agar plates containing Zn for 5 days, followed by 16 hours in Zn-deficient (-Zn) or Zn-excess (10x Zn) as described in Figure 7. Prior to imaging, plants were treated with 100 µM CHX for 4 hours. The mCitrine signal is shown in Fire (LUT) color scale. Scale bars: 25 µm.

## Notes

### Competing Interest Statement

The authors have declared no competing interest.

